# Atypical endo-β-1,4-mannanases are necessary for normal glucomannan synthesis in Arabidopsis

**DOI:** 10.1101/2025.02.21.639209

**Authors:** Aina Kikuchi, Eriko Sato, Yoshihisa Yoshimi, Hironori Takasaki, Naho Nishigaki, Kimie Atsuzawa, Yasuko Kaneko, Masatoshi Yamaguchi, Daisuke Takahashi, Paul Dupree, Toshihisa Kotake

## Abstract

The molecular mechanisms underlying the synthesis of large cell wall polysaccharides in plant cells are not fully understood. Here we report that two atypical endo-β-1,4-mannanases (MANs), which are not secreted and do not degrade glucomannan in the cell wall, play a novel role in glucomannan synthesis. Among the six MANs in Arabidopsis, AtMAN2 and AtMAN5 contain a transmembrane domain at their N-terminal region instead of a signal peptide. Subcellular localization using MAN protein fused with fluorescent protein demonstrated that AtMAN2 localizes to the endomembrane system including the Golgi apparatus in xylem and interfascicular fiber cells. We found that an Arabidopsis *man2 man5* double mutant lost 65% of glucomannan in the cell walls of the inflorescence stem. Immunostaining and immunoelectron microscopic observation also revealed that the *man2 man5* double mutant lost glucomannan in the cell walls to about the same extent as the *csla2 csla9* double mutant, which lacks major glucomannan synthases. Gene complementation experiments showed that the enzymatic activities of AtMAN2 and AtMAN5 are important for the function in synthesis of glucomannan for the cell wall. Arabidopsis possesses another atypical MAN, AtMAN6, with an HDEL retention signal at its C-terminus. However, mutation of AtMAN6 did not affect glucomannan content in the cell walls, suggesting distinct functions for these MANs. This study has identified AtMAN2 and AtMAN5 as novel factors necessary for the normal glucomannan synthesis in Arabidopsis, along with GDP-mannose generating enzymes and CslAs, and suggests that the glucomannan hydrolysis by these MANs contributes to the maintenance of glucomannan synthesis.

## INTRODUCTION

Plant cell wall polysaccharides constitute the largest biomass carbon reservoir on earth. Understanding the mechanism for their synthesis is thus indispensable for their efficient production and utilization (Bar-on et al. 2018). Large polysaccharides are synthesized by glycosyltransferases from building block nucleotide sugars (Verbančič et al. 2018), but the molecular mechanisms are not fully understood. It is well-known that proteins undergo quality control during their synthesis, which prevents protein aggregation and removes unfolded proteins from the endoplasmic reticulum (ER) to be degraded in 26S proteasomes in the cytosol (Mori 2000; Mayer and Bulau 2005). On the other hand, how the aggregation of large cell wall polysaccharides is prevented to maintain the synthesis reaction in the Golgi apparatus is still unknown.

Mannan polysaccharides are cell wall polysaccharides widely distributed in land plants. They commonly possess a backbone composed of β-1,4-mannose (β-1,4-Man) and/or β-1,4-glucose (β-1,4-Glc), while the backbone sequences and decorations are diverse (Scheller and Ulvskov 2010; Voiniciuc 2022; Yoshimi et al. 2025). The backbones of mannan polysaccharides can take on a two-fold structure, in which the sugar residues turn at 180° resulting in a straight ribbon structure suitable for cellulose binding (Yu et al. 2018; 2022). Mannan polysaccharides without decoration tend to become insoluble, probably as a result of intermolecular hydrogen bonding and hydrophobic interactions (Chanzy et al. 1984; Zhou et al. 2018).

The backbones of mannan polysaccharides are synthesized from GDP-Man and GDP-Glc by the action of specific glycosyltransferases, called cellulose-synthase like A (CslA) proteins (Dhugga et al. 2004; Concklin et al. 1999; Liepman et al. 2005). In Arabidopsis (*Arabidopsis thaliana*), CslA2 synthesizes the backbone for β-galactoglucomannan (β-GGM) with alternating β-1,4-Man and β-1,4-Glc residues, whereas ClsA9 makes the backbone for acetylated GGM (AcGGM) with a random sequence of β-1,4-Man and β-1,4-Glc residues (Goubet et al. 2009; Yu et al. 2018; 2022, in this paper, glucomannan is taken to include β-GGM and AcGGM). The reduced glucomannan content in Arabidopsis *vitamin c defective1*, *konjac1*, *csla2,* and *csla9* mutants is an indication of the importance of factors involved in the building block generation and the backbone synthesis reactions (Conklin et al. 1999; 2000; Goubet et al. 2009; Sawake et al. 2015; Nishigaki et al. 2021). These backbones subsequently undergo decoration with α-galactose by mannan α-galactosyltransferase and/or acetyl groups by mannan *O*-acetyltransferase, with which the solubility of glucomannan increases (Reid et al. 2003; Voiniciuc et al. 2015; Zhong et al. 2018; Yoshimi et al. 2025). MANNAN SYNTHESIS-RELATED (MSR) proteins are other factors that have been shown to affect the glucomannan content, although their molecular functions are still obscure (Wang et al. 2013). The identification of additional factors is important for understanding the molecular mechanism of glucomannan synthesis.

Endo-β-1,4-mannanase (MAN) is one of the conserved cell wall enzymes in plants, belonging to the glycoside hydrolase (GH) 5 family (Aspeborg et al. 2021). The physiological importance of MANs has been extensively studied in seed germination and fruit ripening (Nonogaki et al. 2000; Schröder et al. 2004; Belotserkovsky et al. 2007; Iglesias-Fernández et al. 2011). Based on domains, motifs, and sequence similarity, plant MANs can be classified into several types (Fig. 1A) (Wang et al. 2014). Arabidopsis secreted type MANs, AtMAN3 and AtMAN7, hydrolyze cell wall glucomannan to contribute to cadmium tolerance via activation of phytochelatin and glutathione synthesis (Chen et al. 2015; Wu et al. 2023). AtMAN6 is a HDEL-type MAN with a C-terminal HDEL signal, which is a retrograde transport signal back from the Golgi apparatus to the ER (Alvim et al. 2023). AtMAN6 and its poplar (*Populus trichocarpa*) orthologue, PtrMAN6, are specifically localized in vessel cells and presumed to release GGM oligosaccharides as signaling molecules, which negatively regulate secondary cell wall formation in the vessel and interfascicular fiber cells (Zhao et al. 2013; Zhang et al. 2023). On the other hand, AtMAN2 and AtMAN5 are a different type of MAN with a transmembrane (TM) domain instead of a signal peptide in the N-terminal region, designated TM-type MANs here. Recombinant AtMAN2 expressed in *Escherichia coli* shows hydrolytic activity toward mannan polysaccharides and lacks transglycosylation activity (Wang et al. 2015), but its physiological importance and functions remain to be studied.

**Figure 1.**
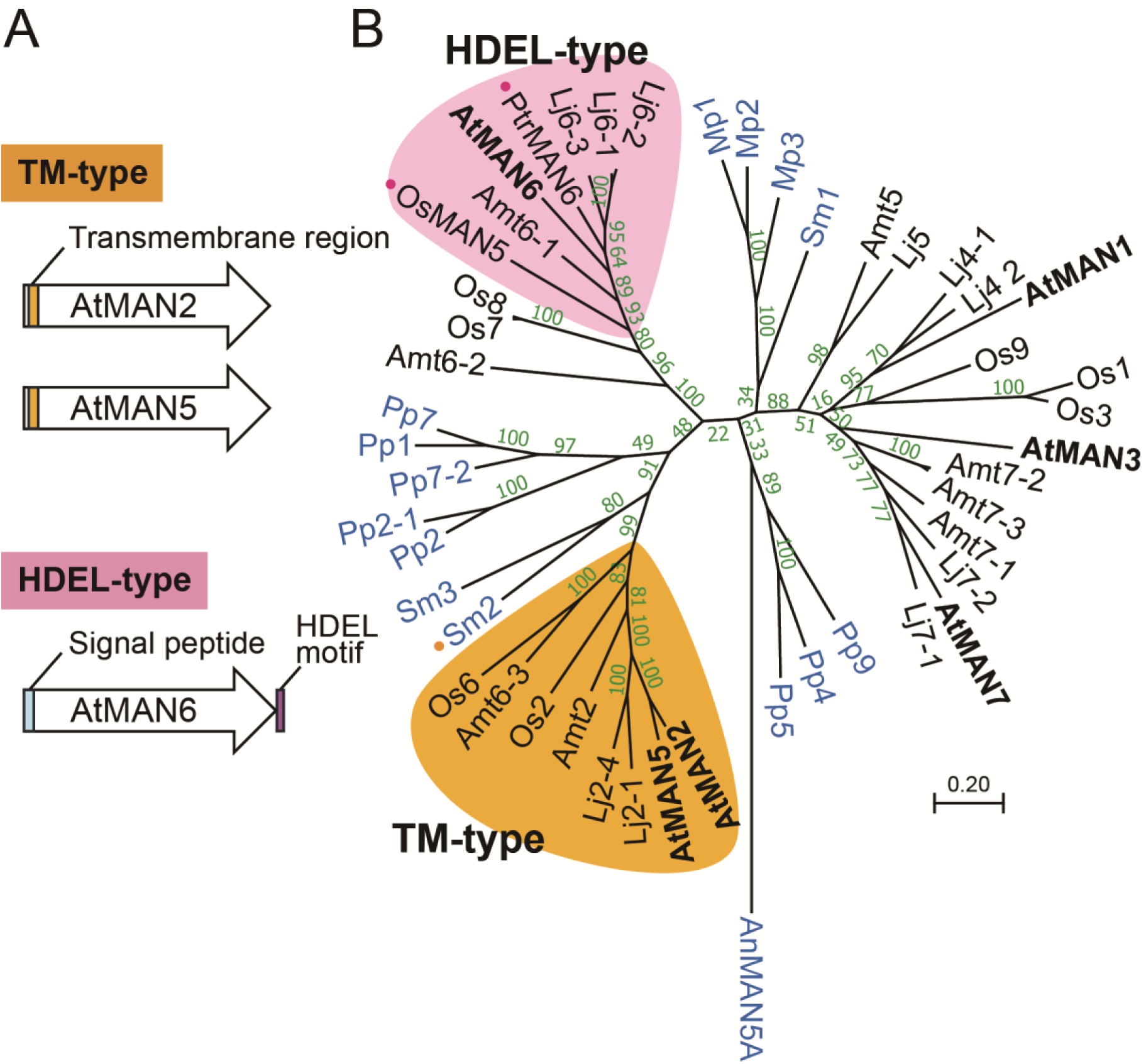
Distribution of TM-type MANs in plants. (A) Schematic models of AtMAN proteins. The signal peptide and TM region were identified with the aid of SignalP 6.0 and TMHMM 2.0 programs, respectively. (B) Phylogenetic tree of plant MANs. Phylogenetic relationships of MANs from Arabidopsis and other plants including *S. moellendorffii*, *P. patens*, and *M. polymorpha* were analyzed with MEGA software (version 11.0). Rice OsMAN5 and poplar PtrMAN6 indicated with pink dots have HHEL and HDDL sequences at the C-terminus instead of HDEL, respectively. SmMAN2 with the orange dot has a TM domain in the N-terminal region that its neighbor, SmMAN3, lacks. MAN sequences from angiosperms are shown in black whereas those from other organisms are in blue. Amt, amborella; Lj, *Lotus japonicus*; Mp, *M. polymorpha*; Os, rice; Pp, *P. patens*; Sm, *S. moellendorffii*. AnMAN5A from *Aspergillus nidulans* was used as the outgroup. Numbers in the nodes represent bootstrap values. The accession numbers and abbreviated names for these sequences are listed in Supplementary Table S2.

This study focused on the importance and functions of the TM-type MANs, AtMAN2 and AtMAN5, comparing them with another atypical MAN, AtMAN6, in glucomannan synthesis. The analysis of Arabidopsis loss-of-function mutants suggested that the hydrolysis of glucomannan by AtMAN2 and AtMAN5 is necessary for normal glucomannan synthesis, where their roles must thus be different from that of AtMAN6. These findings provide new insights into the synthesis of plant cell wall polysaccharides.

## RESULTS

### TM-type MANs are conserved in angiosperms

In Arabidopsis, two TM-type MANs, AtMAN2 (AT2G20680) and AtMAN5 (AT4G28320), are highly related in their amino acid sequences (identity, 83%), while AtMAN6 (AT5G01930) is the sole HDEL-type MAN (Fig. 1A). TM-type MANs can be found in other angiosperms including amborella (*Amborella trichopoda*) and rice (*Oryza sativa*), forming a clade apart from the secreted type MANs in the phylogenetic tree (Fig. 1B). On the other hand, *Physcomitrium patens* (a moss species) and *Marchantia polymorpha* (a liverwort species) lack this type. The HDEL-type MANs also formed a different clade, which did not include MANs from *P. patens* and *M. polymorpha*. The distribution and phylogenetic relationships suggest that the TM-type MANs have functions different from the secreted and the HDEL-type MANs.

### Loss of glucomannan in *man2 man5* double mutant

Because high sequence similarity suggested functional redundancy of AtMAN2 and AtMAN5, an Arabidopsis *man2-1 man5-1* double mutant was generated for functional analysis. The double mutants did not show significant growth defects in the seedlings (Supplementary Fig. S1). In addition, although the double mutants had slightly shorter inflorescence stems than wild type (WT) plants as was also observed for the *csla2-2 csla9-2* double mutants, they did not show severe growth defects (Yu et al. 2022) (Supplementary Fig. S2). Based on the expression patterns (Winter et al. 2007), *AtMAN2* and *AtMAN5* are presumed to function in the inflorescence stem (Supplementary Fig. S3).

To examine the importance of AtMAN2 and AtMAN5 for glucomannan accumulation, we observed the cross sections of the bottom part of inflorescence stems of Arabidopsis *man2-1* single, *man2-2* single, *man5-1* single, and *man2-1 man5-1* double mutants and compared them with WT with LM21 antibody specific to glucomannan (Thorne et al. 2023). Although MANs are hydrolytic enzymes acting on mannan polysaccharides (Wang et al. 2015), the glucomannan signals were, unexpectedly, almost lost in the cell walls of the *man2-1 man5-1* double mutant as they were in the *csla2-2 csla9-2* double mutant, while the signals were observed in the single mutants as they were in the WT, indicating the functional redundancy of AtMAN2 and AtMAN5 (Fig. 2). To examine the functional relationships of AtMAN2 and AtMAN5 with AtMAN6, the cross sections of the *man6-1* single, the *man2-1 man6-1* double, the *man5-1 man6-1* double, the *man2-1 man5-1 man6-1* triple, and the *man2-2 man5-1 man6-1* triple mutants were also observed. The cell wall glucomannan signals were almost lost in the triple mutants as in the *man2-1 man5-1* double mutant but remained in the other double mutants (Fig. 2), suggesting that the AtMAN2 and AtMAN5 contribution to the glucomannan accumulation differs from that of AtMAN6. Xylan, together with cellulose, is a major component of secondary cell walls, but immunostaining with LM10 antibody specific to xylan did not point to compensatory changes in the mutants (Supplementary Fig. S4). The TM-type MANs do not seem to influence the content of other cell wall polysaccharides in the inflorescence stem.

**Figure 2.**
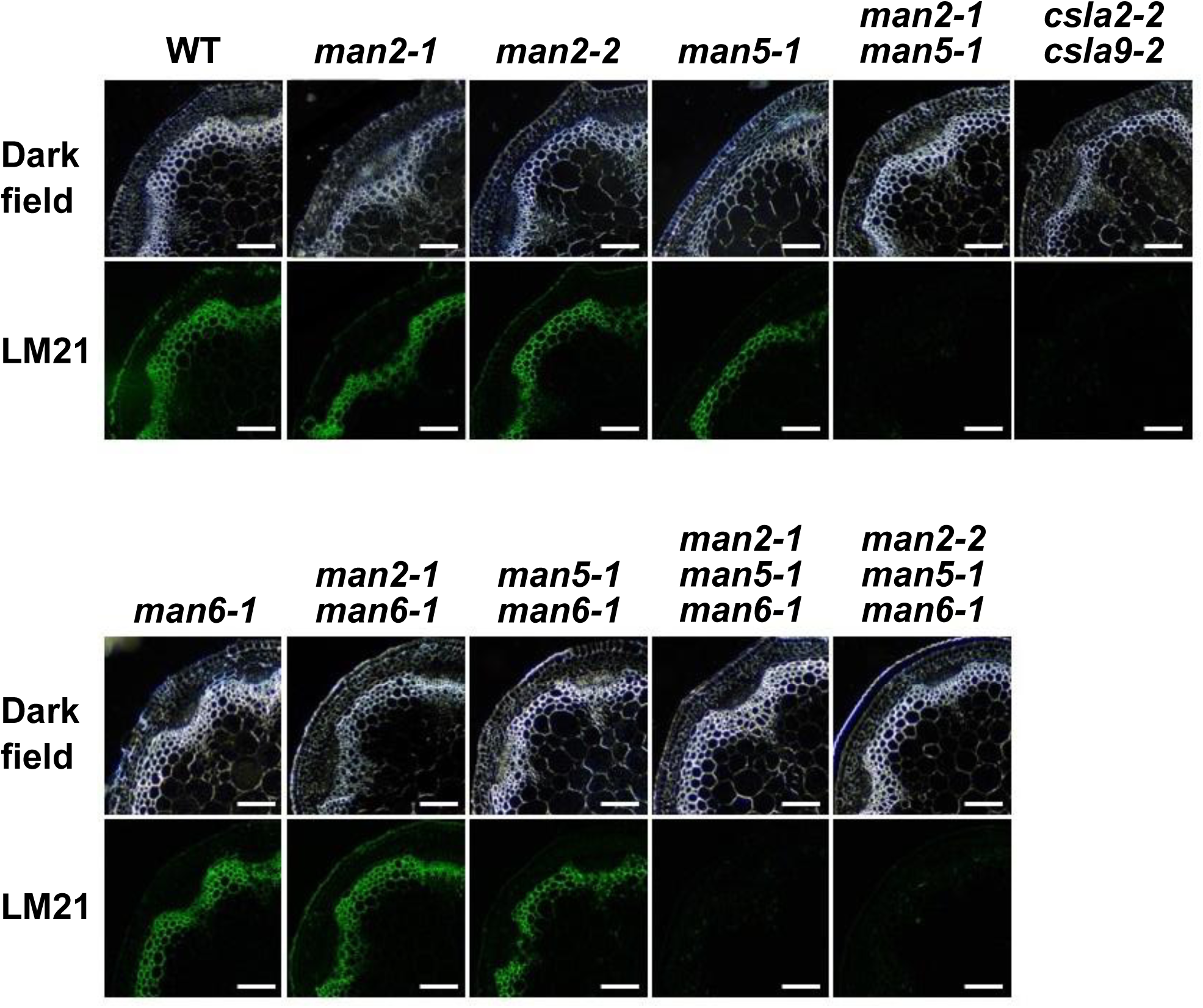
Reduced glucomannan accumulation in the cell walls of *man* mutants. Cross sections from the bottom part of the first inflorescence stem of *man* mutants were compared with WT and the *csla2-2 csla9-2* double mutant. The images for dark field and glucomannan signals with LM21 antibody are shown. The bars indicate 100 μm.

### Drastic decrease of cell wall glucomannan in the double and triple mutants

The changes in the glucomannan accumulation in the inflorescence stems of the *man2-1 man5-1* double and the *man2-1 man5-1 man6-1* and the *man2-2 man5-1 man6-1* triple mutants were further investigated by cell wall fractionation and subsequent sugar composition analysis. The cell walls were fractionated into water-soluble fraction (soluble fraction), alkali-extractable fraction (alkali fraction), and cellulose fraction (Fig. 3A; Supplementary Fig. S5), and the sugar composition of these fractions was determined by high-performance anion-exchange chromatography with pulsed amperometric detection (HPAEC-PAD) (Supplementary Fig. S6). Consistent with the immunostaining of glucomannan with LM21 in the cross sections (Fig. 2), the *man2-1 man5-1* double and the *man2-1 man5-1 man6-1* and *man2-2 man5-1 man6-1* triple mutants lost more than 65% mannose in these cell wall fractions (Fig. 3B). As other hemicelluloses and pectin rarely have mannosyl residues in Arabidopsis (Mohnen 2008; Goubet et al. 2009; Scheller and Ulvskov 2010), this result indicates that the double and triple mutants lost a large part of its glucomannan. It has been reported that AtMAN6 negatively regulates secondary cell wall thickening (Zhang et al. 2023), but neither mannose content nor the sugar content proportion of cell wall fractions was changed in the *man6-1* mutant (Fig. 3).

**Figure 3.**
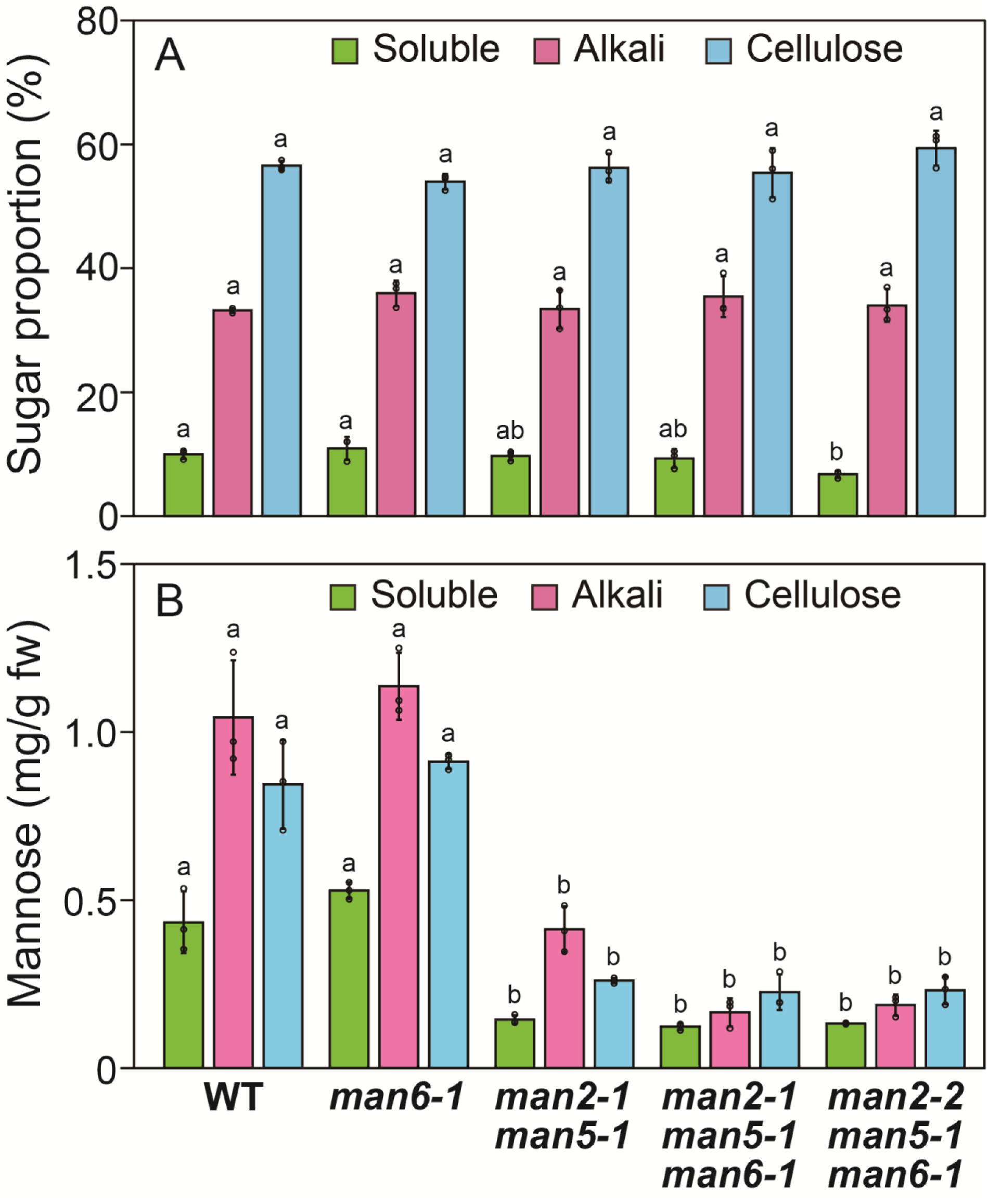
Cell wall fractions and mannose content in *man* mutants. (A) Relative sugar content of cell wall fractions in the *man* mutants. Sugar content of soluble, alkali, and cellulose fractions from the bottom part of inflorescence stems was quantified by the phenol-sulfuric acid method (Supplementary Fig. S4). (B) Mannose content of cell wall fractions. The content was calculated based on the sugar content and the monosaccharide composition of cell wall fractions (Supplementary Fig. S5). The values are shown in weight per gram fresh weight (g fw). Data are mean values with standard deviation (three biological replicates). Different letters indicate significant differences determined with the Tukey HSD test (p < 0.05).

The Arabidopsis inflorescence stem has two different glucomannans, β-GGM and AcGGM, which are synthesized by CslA2 and CslA9, respectively (Goubet et al. 2009; Yu et al. 2018; 2022). The influence of the loss of TM-type MANs on these glucomannans was evaluated by polysaccharide analysis using carbohydrate gel electrophoresis (PACE) (Supplementary Fig. S7). Here, β-GGM and AcGGM included in the stem cell walls were hydrolyzed into oligosaccharides by a microbial MAN, CjMan26A, separated, and detected (Hogg et al. 2001). The *man2-1 man5-1* double and the triple mutants with *man6-1* showed almost no β-Glc-(β-Gal-α-Gal-)β-Man-β-Glc-(α-Gal-)β-Man-β-Glc-Man (GBGAGM) and β-Glc-(β-Gal-α-Gal-)β-Man-β-Glc-Man (GBGM) oligosaccharides from β-GGM and no β-Glc-β-Glc-Man (GGM) and β-Glc-β-Man-Man (GMM) from AcGGM (Supplementary Fig. S7), so they were similar to the *csla2-2 csla9-2* double mutant in this respect (Goubet et al. 2009) (Fig. 4). These results demonstrate that the TM-type MANs are necessary for the normal synthesis of both β-GGM and AcGGM in the inflorescence stem.

**Figure 4.**
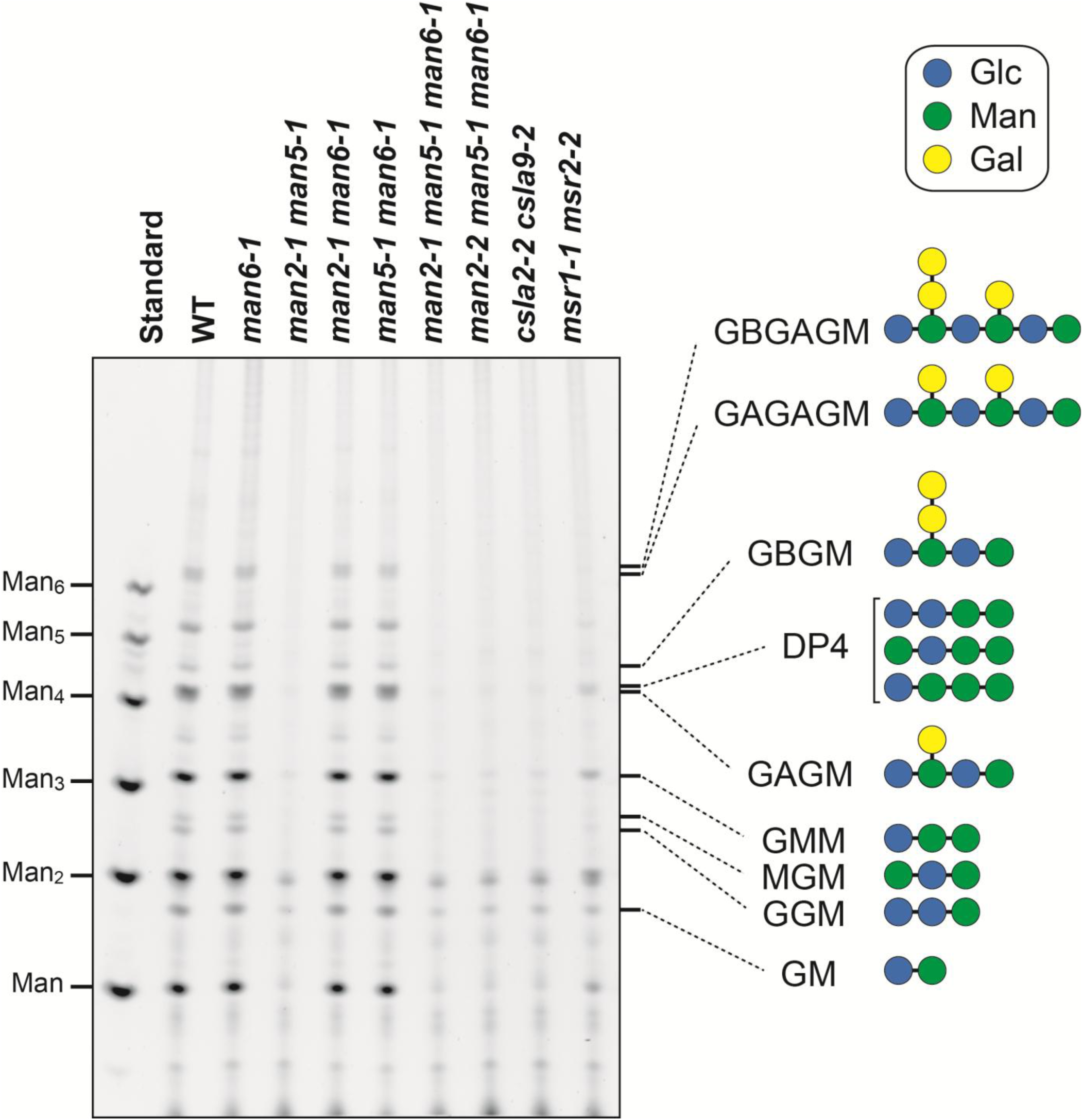
Change in β-GGM and AcGGM. GGMs included in the cell walls of the bottom part of inflorescence stems were hydrolyzed into specific oligosaccharides such as MGM, GBGM, (GA)_2_GM (Supplementary Fig. S6), which were labelled with a fluorescent compound then separated in the gel. Here, *man* mutants were compared with WT, *csla2-1 csla9-1* double mutants known to lack stem glucomannans, and the *msr1-1 msr2-1* double mutant with reduced glucomannan (Goubet et al. 2009; Wang et al. 2013). The sugar symbols follow the international glycan code nomenclature (www.ncbi.nlm.nih.gov/glycans/snfg.html). The oligosaccharide structures are shown on the right. The results for control reactions are shown in Supplementary Fig. S7.

### Retention of glucomannan in the vacuole

β-GGM and AcGGM are synthesized in the Golgi apparatus by CslA2 and CslA9, respectively, then secreted to the cell walls. While the expression levels of these *CslA* genes did not decrease (Supplementary Fig. S8), the glucomannan amount drastically decreased (Figs. 3 and 4). To gain insight into the functions of AtMAN2 and AtMAN5, the glucomannan in the cross sections of *man* mutants was observed at high magnification. Although the *man2-1 man5-1* double and the *man2-1 man5-1 man6-1* triple mutants lost most cell wall glucomannan as shown in Fig. 3, these mutants contained relatively large (>1 μm) particles detected with LM21 antibody inside the xylem and interfascicular fiber cells of the bottom part of inflorescence stems (Fig. 5A). Importantly, glucomannans inside cells have also been observed in Arabidopsis and other plants such as aloe (*Aloe vera*) (Ahl et al. 2019; Handford et al. 2003; Herburger et al. 2022), although these glucomannans have not been identified. Because the particles significantly decreased in the *csla2-2 csla9-2* double mutant, most of these particles can be presumed to be β-GGM and AcGGM that were not secreted to the cell walls. On the other hand, the particles that remained even in the *csla2-2 csla9-2* double mutant may be glucomannan synthesized by different CslAs from CslA2 and Csla9. Although significant differences from WT were not observed, these particles tended to increase rather than decrease in the double and triple mutants (Fig. 5B).

**Figure 5.**
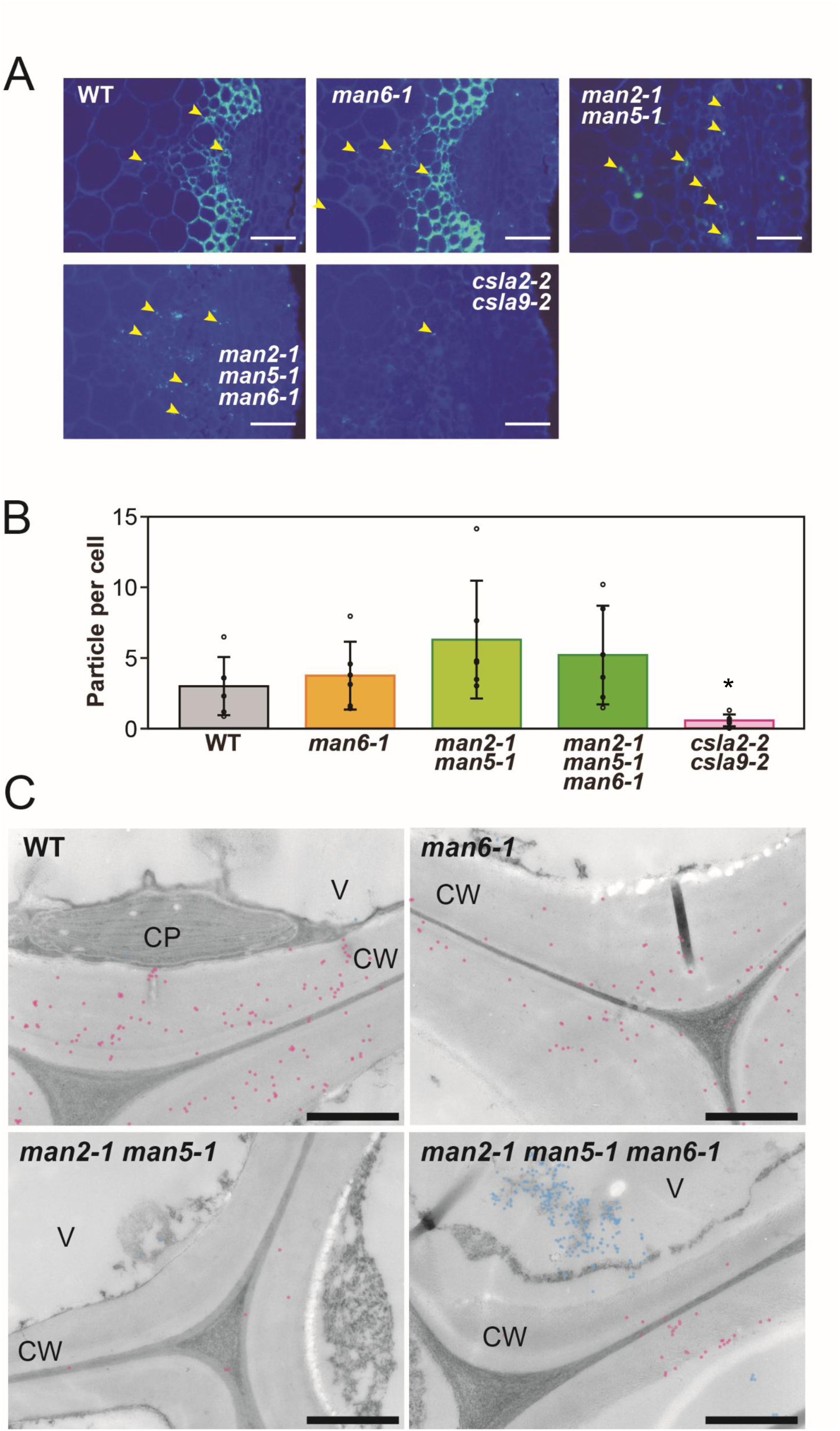
Glucomannan particles inside cells. (*A*) Magnified images of xylem and interfascicular fiber cells. The merged image of calcofluor staining and glucomannan signals with LM21 antibody are shown. The bars indicate 50 μm. Arrowheads indicate relatively large (>1 μm) glucomannan signals found inside cells. (*B*) Numbers of glucomannan particles in the pith, xylem, and interfascicular fiber cells. The values shown are the number of particles per cell. Data are mean values with standard deviation (WT, n□=□6; *man6-1*, n□=□6; *man2-1 man5-1*, n□=□6; *man2-1 man5-1 man6-1*, n□=□6; *csla2-2 csla9-2*, n□=□5). The asterisk indicates significant difference from WT (Student’s t test, *, P < 0.05). (C) Observation by immunoelectron microscopy. Ultrathin sections prepared from the bottom part of the inflorescence stem of WT, *man6-1* single, *man2-1 man5-1* double, and *man2-1 man5-1 man6-1* triple mutants were observed. Glucomannan was detected with LM21 antibody. The gold particles found in the cell walls are shown as red dots and those in the cytoplasm including the vacuole are shown as blue dots. Representative images are shown. CP, chloroplast; CW, cell wall; V, vacuole. The bars indicate 1 µm.

To identify the glucomannan particles retained inside cells, immunoelectron microscopy was conducted. In this experiment, the glucomannans in the ultrathin cross sections from the bottom part of inflorescence stems of the *man2-1 man5-1* double and the *man2-1 man5-1 man6-1* triple mutants were recognized by LM21 as the primary antibody and visualized with a secondary antibody conjugated to gold particles. Consistent with immunostaining and cell wall analysis, there were fewer gold particles in the cell walls of the triple mutant, showing loss of glucomannan in the cell walls. Relatively large clusters (>1 μm) of gold particles were found in the vacuole in the triple mutant (Fig. 5C). These observations suggest that the glucomannan particles are formed and retained in the vacuole in the absence of the TM-type MANs, AtMAN2 and AtMAN5.

### Tissue and subcellular localization of AtMAN2 and AtMAN6 differ

The tissue and subcellular localization of AtMAN2 and AtMAN5 was investigated using transgenic Arabidopsis harboring *pMAN2::MAN2TM-mNeon* or *pMAN5::MAN5TM-mNeon* in which the fluorescent protein mNeonGreen sequence was inserted right behind the TM domain in the genomic sequences (Supplementary Fig. S9). In this experiment, the gene constructs were introduced into the *man2-1 man5-1 man6-1* or the *man2-2 man5-1 man6-1* triple mutants. Immunostaining of the cross sections of the bottom part of inflorescence stems showed that AtMAN2TM-mNeon and AtMAN5TM-mNeon at least partially restored impaired glucomannan accumulation in the triple mutants (Supplementary Fig. S9). For microscopic observation, the top part of the inflorescence stem was chosen because strong mNeonGreen signals were found there. The AtMAN2TM-mNeon signal was widely observed in the xylem and interfascicular fiber cells of inflorescence stems (Supplementary Fig. S10). The AtMAN2TM-mNeon signal was not found in the cell walls but localized in the compartments (Fig. 6A-E), suggesting that AtMAN2 functions in the endomembrane including the Golgi apparatus. The AtMAN5TM-mNeon signal was also found inside cells (Fig. 6F-J), although the localization was not clear because of low signal intensity.

**Figure 6.**
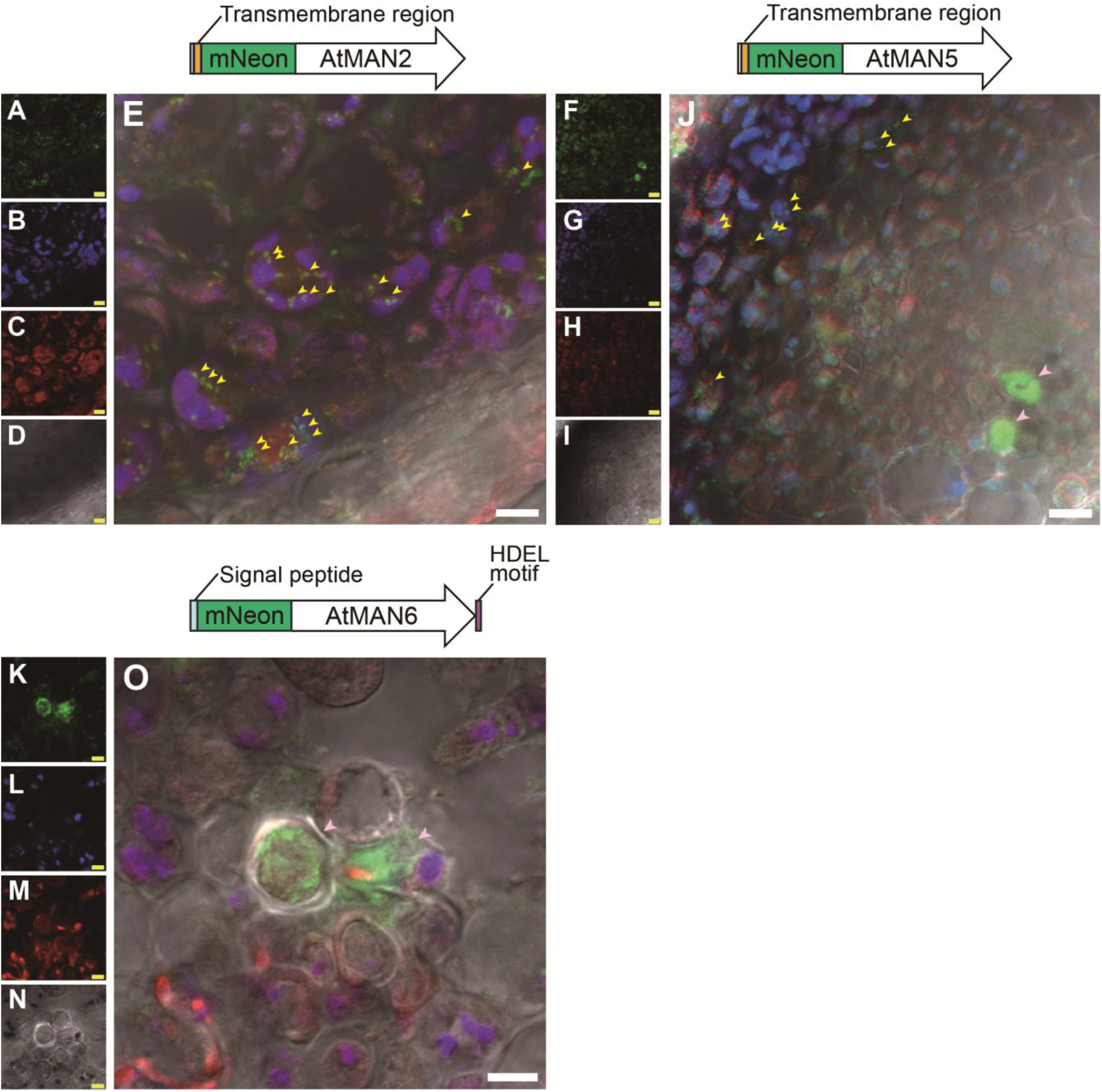
Subcellular localization of MAN-mNeon proteins. Xylem and interfascicular fiber cells expressing AtMAN2TM-mNeon (A-E) and AtMAN5TM-mNeon (F-J), and vessels expressing AtMAN6signal-mNeon (K-O) in the top part of the inflorescence stem were observed with a confocal microscope. Images for mNeonGreen (A, F, K), autofluorescence (B, G, L), FM 4-64FX staining (C, H, M), and transmitted photomultiplier tube (D, I, N) were obtained and merged (E, J, O). The bars indicate 5□μm. Schematic diagrams for each mNeonGreen (mNeon)-MAN protein are shown above the merged images (E, J, O). The representative mNeonGreen signals in the endomembrane system including the Golgi apparatus are indicated by yellow arrowheads and those in vessels with by pink arrowheads. The gene constructs and the complementation of the triple mutants with *pMAN2:MAN2TM-mNeon* and *pMAN5:MAN5TM-mNeon* are shown in Supplementary Fig. S9. Images of the tissue localization of these MAN-mNeon proteins and longitudinal sections for AtMAN6signal-mNeon are shown in Supplementary Fig. S10.

To compare, we also generated *pMAN6::MAN6signal-mNeon* in which the mNeonGreen sequence was inserted after the signal peptide into the genomic *AtMAN6* sequence (Supplementary Fig. S9). Contrasting with AtMAN2TM-mNeon signals, the AtMAN6signal fluorescence was limited to developing vessels (Supplementary Fig. S10). The fluorescence was spread in the vessel cells (Fig. 6K-O), indicating its localization in the ER or the vacuole. Previously it was suggested that AtMAN6 localizes on the plasma membrane (PM), but this was using a C-terminal fusion of AtMAN6 with green fluorescent protein, which might have affected the localization. But in that study, as in this, AtMAN2 and AtMAN6 were found in substantially different locations, indicating that they have different molecular functions (Zhang et al. 2023).

### The enzymatic activities of AtMAN2 and AtMAN5 is required

To determine the importance of the enzymatic activities of AtMAN2 and AtMAN5, we performed gene complementation experiments on the *man2-1 man5-1* double mutant using point-mutated genomic *AtMAN2* and *AtMAN5* genes in which the catalytic residues were replaced with Ala (E215A and E335A in AtMAN2; E214A and E334A in AtMAN5). The mutated *AtMAN2* and *AtMAN5* genes failed to restore the glucomannan accumulation detected with the LM21 antibody in the inflorescence stem of the double mutant (Supplementary Figs. S9, S11). This result suggests that for proper functioning of the glucomannan synthesis in the Golgi apparatus, the enzymatic activity of these MANs is necessary.

## DISCUSSION

### MANs are newly recognized as necessary for normal glucomannan synthesis

The present study has identified AtMAN2 and AtMAN5 as factors necessary for normal glucomannan synthesis in Arabidopsis, to work along with VTC1 and KJC1 generating GDP-Man, CslAs catalyzing the glucomannan backbone synthesis, and MSRs (Conklin et al. 1999; Goubet et al. 2009; Wang et al. 2013; Yu et al. 2014; Sawake et al. 2015; Nishigaki et al. 2021). Based on the distribution of orthologues, it is likely that many angiosperms utilize TM-type MANs in glucomannan synthesis.

Because the expression levels of *CslA2* and *CslA9* did not decrease in the *man2-1 man5-1* double and the *man2-1 man5-1 man6-1* triple mutants (Supplementary Fig. S8), we infer that the decreased glucomannan in the mutants was not caused by changes in the expression of *CslA* genes and that AtMAN2 and AtMAN5 contribute to glucomannan synthesis through their enzymatic activity. The requirement of polysaccharide degrading enzymes for polysaccharide synthesis presents an apparent paradox. Yet this is not the first example: among other plant glycoside hydrolases, the membrane-associated GH9 endo-β-1,4-glucanase KORRIGAN1 (KOR1) is well known for its role in cellulose synthesis (Nicol et al. 1996). The Arabidopsis *kor1* mutant suffers severe defects in the synthesis of cellulose, demonstrating the importance of this protein. The sea squirt *Ciona intestinalis* and bacteria such as *Agrobacterium tumefaciens* and *Acetobacter xylinum* also require endo-β-1,4-glucanases for their cellulose synthesis. These enzymes belong to the GH6 and 8 families and are thus distinct from KOR1 (Standal et al. 1994; Matthysse et al. 1995; Li et al. 2023). These observations imply that hydrolytic activity of endo-β-1,4-glucanases is important to facilitate cellulose synthesis. The present study describes a second instance of a version of this paradox in polysaccharide synthesis. We suggest that this paradox may represent a general mechanism to prevent the aggregation of newly synthesized cell wall polysaccharides in plant cells.

### Not all CslAs require TM-type MANs for glucomannan synthesis

The TM-type MANs are not necessarily required for all glucomannan synthesis by CslAs in plants. In the present study, the alkali fraction from the inflorescence stems of *man2-1 man5-1* double mutant still contained 30% of mannose compared with WT (Fig. 3B). Because *N*-glycosylated proteins with mannosyl residues would not have been collected in the alkali fraction, there seems to be mannan polysaccharide synthesized independently from the functions of AtMAN2 and AtMAN5. This unknown mannan polysaccharide would have to be resistant to hydrolysis by CjMan26A used for PACE.

Many green algae and bryophytes such as *P. patens* and *M. polymorpha* have mannan polysaccharides as major cell wall components (Kolkas et al. 2023; Chernova et al. 2024), although they lack genes orthologous to *AtMAN2* and *AtMAN5* (Fig, 1B). To understand the molecular mechanism for the glucomannan synthesis involving TM-type MAN, future research should try to clarify which types of mannan polysaccharides require these MANs.

### Putative function of AtMAN2 and AtMAN5 in quality control

AtMAN2TM-mNeon localized in the endomembrane including the Golgi apparatus in the inflorescence stem (Fig. 6). In the gene complementation experiments, the genomic *AtMAN2* and *AtMAN5* genes with mutations at catalytic residues failed to restore cell wall glucomannan in the *man2 man5* double mutant (Supplementary Fig. S11). Recently, AtMAN2 and AtMAN5 were also shown to increase soluble mannan polysaccharides in Pichia yeast (*Pichia pastoris*) expressing Arabidopsis *CslA2* (Jacobson et al. 2025). It has been reported that AtMAN2 lacks transglycosylation activity (Wang et al. 2015). Based on these facts, we suggest that glucomannan undergoes hydrolysis by AtMAN2 and AtMAN5 after the synthesis in the Golgi apparatus. The drastic decrease in the glucomannan content in the double and triple mutants implies that the continuous synthesis reaction by CslAs is not maintained in these mutants. The alternative, that only secretion is impaired in the mutants, while the glucomannan synthesis by CslA2 and CslA9 is maintained, does not seem likely, because in this case the glucomannan would have been collected in one of the fractions (Fig. 3B). We cannot specify the molecular functions of these MANs in detail, but it is conceivable that large β-GGM and AcGGM molecules synthesized by CslA2 and CslA9 before decoration easily aggregate during synthesis, which may cause the termination of the chain elongation reaction by these CslAs. This problem may be alleviated through removal by partial hydrolysis with these MANs as polysaccharides generally become more soluble through hydrolysis into shorter chains. These reactions to prevent aggregate formation and to remove aggregates in the Golgi apparatus can be considered analogous to the protein quality control in the ER (Mori 2000; Oda et al. 2003). It is also probable that AtMAN2 and AtMAN5 facilitate the release of glucomannan from CslA proteins by partial hydrolysis during synthesis, enabling its proper loading into the secretion pathway. The impairment of this process may also negatively affect glucomannan synthesis by CslAs.

In xyloglucan, the importance of galactose-side chains to avoid aggregation has been shown (Tamura et al. 2005; Kong et al. 2015; Hoffmann and McFarlane 2024). The Arabidopsis *murus3* mutant, which lacks galactose-side chains on its xyloglucan, suffers intracellular aggregation of xyloglucan in the roots, resulting in growth defects. It is probable that such intracellular aggregation was avoided through the termination of the synthesis reaction in the *man2-1 man5-1* double mutant. Clarifying this mechanism will presumably require the identification of as yet unknown factors involved in this reaction.

## MATERIALS AND METHODS

### Arabidopsis mutants and growth conditions

*Arabidopsis thaliana* ecotype Col-0 was used in this study. The T-DNA insertion lines SALK_0383344 (*man2-1*), SALK_016450 (*man2-2*), SALK_059047 (*man5-1*), and SALK_129977 (*man6-1*) were provided by the Arabidopsis Biological Resource Center. The double and triple mutants were generated by crosses between these mutants. The genotypes of the mutants were determined using specific primers (Supplementary Fig. S12 and Table S1). The *csla2-2 csla9-2* and *msr1-1 msr2-2* double mutants had been obtained in our previous studies (Goubet et al. 2007; 2009; Wang et al. 2013). Arabidopsis seeds were sterilized and sown on Murashige and Skoog (MS) medium in 0.8% (w/v) agar. After stratification at 4 °C in the dark for 2 days, the seeds were germinated and grown under continuous light (cold cathode fluorescent lamp; Biomedical Science) at 23 °C for 10 days, then on rockwool fiber for 4 weeks under continuous light at 23 °C.

For the gene complementation experiments, genomic DNA fragments for *AtMAN2* and *AtMAN5* were amplified by PCR and subcloned into the binary vector pBGGN (Inplanta Innovation, Yokohama, Japan). The point mutations at the catalytic sites were introduced by PCR with specific primers (Supplementary Table S1). The *man2-1 man5-1* double mutant was transformed with mutated AtMAN2/pBGGN, WT AtMAN2/pBGGN, mutated AtMAN5/pBGGN, and WT AtMAN5/pBGGN by an Agrobacterium-mediated (*Rhizobium radiobacter*, EHA105 strain) method (Clough and Bent 1998).

### Phylogenetic analysis

The amino acid sequences of MANs from various plants were collected using Basic Local Alignment Search Tool in NCBI (https://blast.ncbi.nlm.nih.gov/Blast.cgi) (Supplementary Table S2). The signal peptide and TM region were detected using the SignalP 6.0 and TMHMM-2.0 programs, respectively (Teufel et al. 2022; Krogh et al. 2001; Moller et al. 2001). The alignments of amino acid sequences were performed by the MUSCLE program in MEGA software (version 11; Tamura et al. 2021) with default parameters. The phylogenetic trees were generated using the maximum likelihood method with 100 bootstraps.

### Cell wall fractionation and sugar composition analysis

The bottom part of first inflorescence stem 2 weeks after bolting (approximately 50 mg) was homogenized with mortar and pestle in water (Supplementary Fig. S4). After centrifugation at 10,000 x g at 4°C for 5 min, the supernatant was collected as the soluble fraction. The precipitate was suspended in 0.4 mL of 4 M potassium hydroxide (KOH) containing 0.04% (w/v) sodium borohydride at 25°C. After centrifugation at 10,000 x g at 4°C for 5 min, the supernatant was collected. This extraction was repeated. The combined supernatant (0.8 mL) was neutralized with 0.4 mL of glacial acetic acid and collected as the alkali fraction. The precipitate was washed with 98% (v/v) ethanol, 2% (v/v) acetic acid twice, lyophilized for 24 h, hydrolyzed with 72% (v/v) sulfuric acid at 4 °C for 1 h and 8% (v/v) sulfuric acid at 100 °C for 4 h, and then collected as the cellulose fraction. Total sugar composition and quantity of the above fractions were determined using the phenol sulfuric acid procedure (Dubois et al. 1956).

For sugar composition analysis, the alkali fraction was dialyzed against water at 4 °C for 2 days. The soluble and alkali fractions were hydrolyzed into monosaccharides with 2 M trifluoroacetic acid (TFA) at 121°C for 1 h, then lyophilized to remove TFA. The cellulose fraction after the hydrolysis with sulfuric acid was neutralized with barium carbonate and desalted with Dowex beads (Dowex 50WX8, Dow Chemical). Monosaccharide composition was determined by HPAEC-PAD using a Dionex ICS-5000+ liquid chromatograph (Thermo Fisher Scientific) fitted with a CarboPac PA-1 column (Thermo Fisher Scientific) and a pulsed amperometric detector (Thermo Fisher Scientific) as described previously (Sawake et al. 2015).

### PACE

Alcohol-insoluble residue (AIR) was prepared from the bottom part of Arabidopsis inflorescence stems (fresh weight, 100 mg) according to Yu et al. (2018). Glucomannans were extracted from the AIR with 1 mL of 4 M KOH solution by shaking at 1,000 rpm at room temperature for an hour. After centrifugation at 15,000 x g for 15 minutes, the supernatant was collected. To collect remaining glucomannans, the pellet was suspended in 1 mL of water. After centrifugation, the supernatant was collected. The supernatants were combined and applied to a PD-10 column (Cytiva) to exchange the solvent with 50 mM ammonium acetate buffer (pH 6.0). The eluted fraction was used as the KOH fraction.

Glucomannans in the KOH fraction were hydrolyzed into specific oligosaccharides with a commercial endo-β-1,4-mannanase, CjMan26A (1 unit, E-BMACJ, Megazyme), from *Cellvibrio japonicus* in 50 mM ammonium acetate buffer (pH 6.0) at 37 °C overnight (Supplementary Fig. S7) (Hogg et al. 2001). After the reaction, the enzyme was denatured by heating at 100 °C for 15 minutes. The oligosaccharides were derivatized with 8-aminonapthalene-1,3,6-trisulphonic acid, resuspended in urea, and separated by electrophoresis as described (Goubet et al. 2009; Yoshimi et al. 2025). Gels were visualized using a G-Box (Syngene, Cambridge, UK) with a transilluminator with long-wave tubes emitting at 365 nm and a shortpass (500 to 600 nm) detection filter. Mannose and β-1,4-mannooligosaccharides (Man_2-6_, Megazyme) were used as standards.

### Immunostaining of glucomannan in the mutants

The glucomannan and xylan in the inflorescence stems were observed using specific antibodies. The bottom part of inflorescence stems (the lowest one-sixth portion) was fixed with 4% (v/v) paraformaldehyde and 0.6% (v/v) glutaraldehyde at 4 °C for 3 h. The fixed samples were dehydrated through a graded ethanol series, embedded in Technovit 7100 (Nisshin EM, Tokyo, Japan) using a graded series of ethanol and Technovit 7100 mixtures, and solidified with Technovit hardener (Nisshin EM) at 60 °C for 12 h. Sections were sliced to a thickness of 10 μm using a microtome (RM2125RT; Leica, Tokyo, Japan) equipped with a disposable steel blade (C35; Feather, Osaka, Japan).

To remove acetyl groups from polysaccharides, the cross-sections were treated with 50 mM sodium hydroxide at 25 °C for 30 min. The sections were then washed with phosphate-buffered saline (PBS) containing 137 mM NaCl, 10 mM disodium hydrogenphosphate, 2.7 mM KCl, and 1.8 mM potassium dihydrogenphosphate (pH 7.4) and then blocked in PBS containing 0.3% (w/v) blocking milk (Sigma-Aldrich, Tokyo, Japan) for 30 min. Then, the sections were incubated for 1.5 h with antibodies against glucomannan (LM21, Kerafast, Shirley, MA; dilution 1:5) or xylan (LM10, Kerafast; dilution 1:5) in PBS with blocking milk. After washing three times with PBS, the sections were incubated for 1 h with Alexa Fluor 488-conjugated anti-mouse IgM for LM21 (abcam, Cambridge, UK; dilution 1:100) or Alexa Fluor 488-conjugated anti-mouse IgG for LM10 (abcam; dilution 1:100) in PBS with bovine serum albumin (Thorne et al. 2023). The sections were washed three additional times with PBS. The cross sections were imaged with a microscope (ECLIPSE Ci-L Plus, Nikon, Japan) at 488 nm. The dark field image was also acquired simultaneously.

### Immunoelectron microscopy

Immunoelectron microscopy to detect the glucomannan localization was performed according to Atsuzawa et al. (2010). The bottom part (the lowest one-sixth portion) and upper part (the highest one-sixth portion) of the inflorescence stems of WT, the *man6-1* single, the *man2-1 man5-1* double, and the *man2-1 man5-1 man6-1* triple mutants were fixed with 2% (v/v) glutaraldehyde in 50 mM potassium phosphate (pH 6.8) at room temperature for 3 h. After washing with 100 mM lysine in 50 mM potassium phosphate (pH 6.8) the samples were dehydrated in a graded ethanol series. The tissues were embedded in LR White resin (Electron Microscopy Sciences, USA) and polymerized by heating at 60 °C. Ultrathin sections of 100 nm thickness were cut on an UltracutN ultramicrotome (Nissei) and collected on nickel grids coated with polyvinyl formvar membrane.

Ultra-thin sections were immersed in 50 mM sodium hydroxide at room temperature for 30 min and washed with Tris-HCl buffer (TBS: 50 mM Tris-HCl (pH 7.5), 150 mM sodium chloride) containing 0.05% Triton X-100 (TBS/Tx). They were treated with blocking solution (10 mg/ml bovine serum albumin in TBS) at room temperature for 60 min, and they were incubated with LM21 antibody at 1:500 dilution at 4 °C overnight. After washing with TBS/Tx, the sections were incubated with a solution of 10 nm gold particles conjugated with goat anti-rat IgG (BBI solutions, UK) as the secondary antibody at a 1:30 dilution for 1 h at room temperature and washed with TBS/Tx. Specimens were observed with a transmission electron microscope (H-7500, Hitachi, Japan) at an accelerating voltage of 80 kV after staining with uranyl acetate and lead citrate.

### Transgenic plants expressing MAN-mNeonGreen

For the generation of transgenic Arabidopsis harboring *pMAN2::MAN2TM-mNeon*, *pMAN5::MAN5TM-mNeon*, or *pMAN6::MAN6signal-mNeon*, the DNA fragments of genomic *AtMAN2*, *AtMAN5*, and *AtMAN6* and mNeonGreen were amplified from Arabidopsis genomic DNA and template mNeonGreen (Addgene, #189976), respectively, using the primer sets listed in Supplementary Table S1. The DNA fragments were assembled and subcloned into the binary vector pBGGN with DNA assembly kit (Gibson) to yield the constructs, *pMAN2::MAN2TM-mNeon/pBGGN*, *pMAN5::MAN5TM-mNeon/pBGGN* and *pMAN6::MAN6signal-mNeon/pBGGN* (Supplementary Fig. S9). These constructs were introduced into the *man2-1 man5-1 man6-1* or the *man2-2 man5-1 man6-1* triple mutants by an Agrobacterium-mediated (*Rhizobium radiobacter*, EHA105 strain) method (Clough and Bent 1998).

### Observation of tissue and subcellular localization of MAN-mNeonGreen proteins

Cross sections were excised from the top part of the inflorescence stems of *pMAN2::MAN2signal-mNeon* (line #3), *pMAN2::MAN2signal-mNeon* (line #4), and *pMAN6::MAN2signal-mNeon* (line #3) plants and stained with 6.34 μM fixable analog of N-(3-triethylammoniumpropyl)-4-(6-(4-(diethylamino) phenyl) hexatrienyl) pyridinium dibromide, FM 4-64FX (ThermoFisher). The confocal images were obtained using a Zeiss LSM800 (Zeiss, Jena, Germany) with a Plan-Apochromat 63×/1.4 oil immersion objective lens. FM 4-64FX was excited at 561 nm and detected between 400 and 645 nm. Fluorescence signals from mNeonGreen were detected after excitation at 488 nm and detected at 410-546 nm. Autofluorescence was excited at 488 nm and detected at 550-700 nm. Transmitted photomultiplier tube (T-PMT) images were also captured during confocal imaging.

## Author Contributions

T.K. and P.D. designed the research. A.K., E.S., Y.Y., H.T., N.N., K.A., Y.K., D.T., and T.K. performed research. E.S., Y.Y., D.T., Y.K., and T.K. analyzed the data. H.T., Y.K., D.T., P.D., and T.K. wrote the manuscript.

## Acknowledgments

We thank Dr. Kiminori Toyooka from RIKEN and Prof. Masatsugu Toyota and Dr. Hiraku Suda from Saitama University for valuable discussions. This work was supported by MEXT KAKENHI Grant-in-Aid for Scientific Research to T. Kotake [no. 18H05495], Grant-in-Aid for Scientific Research to T. Kotake [no. 23H04302 and no. 23H02134] and to D. Takahashi [no. 23K05144 and no. 24KK0273], the ERC Advanced Grant EVOCATE to P. Dupree funded by the United Kingdom Research and Innovation (UKRI) grant number EP/X027120/1 (www.ukri.org) and by the grant OpenPlant (UKRI BBSRC, BB/L014130/1) to P. Dupree, and a Broodbank Research Fellowship to Y. Yoshimi (reference no. PD16178).

## Supplementary Data

**Supplementary Table S1.** Supplementary Table S1. Primers used for genotyping, plasmid construction, point mutations, and RT-PCR.

**Supplementary Table S2.** List of MAN sequences in the phylogenetic analysis.

**Supplementary Fig. S1.** Growth phenotypes of seedlings of *man* mutants

**Supplementary Fig. S2.** Growth phenotypes in the inflorescence stems of *man* mutants

**Supplementary Fig. S3.** Expression patterns of *AtMAN2*, *AtMAN5*, and *AtMAN6* genes.

**Supplementary Fig. S4.** Xylan accumulation in *man* mutants.

**Supplementary Fig. S5** Cell wall fractionation from inflorescence stems.

**Supplementary Fig. S6.** Monosaccharide composition of the soluble, alkali, and cellulose fractions.

**Supplementary Fig. S7.** Oligosaccharides detected by PACE.

**Supplementary Fig. S8.** Expression levels of *CslA2* and *CslA9* in *man* mutants.

**Supplementary Fig. S9.** Gene constructs and complementation of *man* mutants.

**Supplementary Fig. S10.** Tissue localization of MAN-mNeon proteins.

**Supplementary Fig. S11.** Gene complementation experiment with mutated genomic *AtMAN2* and *AtMAN5* genes.

**Supplementary Fig. S12.** T-DNA insertion sites and genotyping of *man* mutants.

