## Supplementary material for "Atypical endo-β-1,4-mannanases are necessary for normal glucomannan synthesis in Arabidopsis": Sup. Tables

**Supplementary Table S1. Primers used for genotyping, plasmid construction, point mutations, and RT-PCR.**

| | |
|---|---|
| Genotyping | |
| MAN2_genotype_F | 5’-CCTCTGTTAAGAACTGTGTG-3’ |
| MAN2_genotype_R | 5’-CCAAAGTCATCGTTGAAACC-3’ |
| MAN5_genotype_F3 | 5’-GCTCACAGTTAATCCAGAACAG-3’ |
| MAN5_genotype_R3 | 5’-GTCCTTGTTCTGGTTTGATAGG-3’ |
| MAN6_genotype_F | 5’-GAAGTGTTCCAACAAGCCTCTG-3’ |
| MAN6_genotype_R | 5’-GAGTTTTCGCTGAGATGGATGG-3’ |
| LBb1 | 5’-GCGTGGACCGCTTGCTGCAACT-3’ |
| *pMAN:MAN-mNeon* construct | |
| pBGGN_H_gMAN2_F | 5’-CGCATCTACTATATGTTTGAAGCTTCACGAAAGCTCTAAAAAATTCG-3’ |
| gMAN2TM_B_mNeon_R | 5’-CCATGGATCCGAGACCGAACCACAAATCTCC-3’ |
| gMAN2TM_B_mNeon_F | 5’-GTTCGGTCTCGGATCCATGGTGAGCAAGGGCGAG-3’ |
| mNeon_gMAN2_R_1 | 5’-GCCCTCCGTTTTAGATCCCTTGTACAGCTCGTCCATGC-3’ |
| mNeon_gMAN2_F_2 | 5’-TACAAGGGATCTAAAACGGAGGGCGAGTTAGCATTC-3’ |
| gMAN2_E_pBGGN_R | 5’-TGTAAAACGACGGCCAGTGAATTCGATGACTTAGAGACGCAATC-3’ |
| pBGGN_H_gMAN5_F | 5’-CGCATCTACTATATGTTTGAAGCTTCTGAAACCTGAACTATAATATGGG-3’ |
| gMAN5TM_B_mNeon_R | 5’-CCATGGATCCCAACCACAAATCTCGAAATG-3’ |
| gMAN5TM_B_mNeon_F | 5’-TTTGTGGTTGGGATCCATGGTGAGCAAGGGCGAG-3’ |
| mNeon_gMAN5_R_1 | 5’-CCCTTTGTGATTAGATCCCTTGTACAGCTCGTCCATGC-3’ |
| mNeon_gMAN5_F_2 | 5’-TACAAGGGATCTAATCACAAAGGGAAAGCAAAGTTAGGG-3’ |
| gMAN5_E_pBGGN_R | 5’-TGTAAAACGACGGCCAGTGAATTCAACGAGGTCATCTCCAAGACTG-3’ |
| pBGGN_H_gMAN6_F | 5’-CGCATCTACTATATGTTTGAAGCTTAGAAACTCATGAAGATATGGAAA-3’ |
| gMAN6signal_B_mNeon_R | 5’-CCATGGATCCTGCGAGAGCTCTGTTTTGAGTCAG-3’ |
| gMAN6signal_B_mNeon_F | 5’-AGCTCTCGCAGGATCCATGGTGAGCAAGGGCGAG-3’ |
| mNeon_gMAN6_R_1 | 5’-CGCTGTCCAAGTCAGATCCCTTGTACAGCTCGTCCATGC-3’ |
| mNeon_gMAN6_F_2 | 5’-TACAAGGGATCTGACTTGGACAGCGAGTCCCATG-3’ |
| gMAN6_E_pBGGN_R_2 | 5’-TGTAAAACGACGGCCAGTGAATTCTCTGATTTAAATTCTTATGATG-3’ |
| *Point mutations* | |
| MAN2_E215A_F | 5’-TTTGCATGGGCGTTGATAAACGAGCCTCGCTGC-3’ |
| MAN2_E215A_R | 5’-GTTTATCAACGCCCATGCAAAAATTGTTGGGTCG-3’ |
| MAN2_E335A_F | 5’-CTGTTCACCGCGTTTGGACTCTCGAATCTGAACAAG-3’ |
| MAN2_E335A_R | 5’-GAGTCCAAACGCGGTGAACAGAACAGGCTTCTTC-3’ |
| MAN5_E214A_F | 5’-TTCGCTTGGGCGTTGATAAACGAGCCTCGATG-3’ |
| MAN5_E214A_R | 5’-GTTTATCAACGCCCAAGCGAAAATCGTAGGATC-3’ |
| MAN5_E334A_F | 5’-CTCTTCACAGCGTTTGGCCTATCAAACCAGAAC-3’ |
| MAN5_E334A_R | 5’-TAGGCCAAACGCTGTGAAGAGAACTGGCTTCTTTAG-3’ |
| *RT-PCR* | |
| CslA2_RTP_F | 5’-GAGGAGAAGCTCCAAGGGTTC-3’ |
| CslA2_RTP_R | 5’-TGTAGCTCACGCAGAATCTTGA-3’ |
| CslA9_RTP_F | 5’-AACAGCACAGGTGGTCATGT-3’ |
| CslA9_RTP_R | 5’-GGAATGTAAACCGCTCCCCA-3’ |
| ACT2_RTP_F | 5’-TCCCTCAGCACATTCCAGCAGAT-3’ |
| ACT2_RTP_R | 5’-AACGATTCCTGGACCTGCCTCAT-3’ |
| GAPDH_RTP_F | 5’-TTGGTGACAACAGGTCAAGCA-3’ |
| GAPDH_RTP_R | 5’-AAACTTGTCGCTCAATGCAATC-3’ |

**Supplementary Table S2. Abbreviated names and accession numbers**

| Abbreviated name | Source species | Accesion number or gene locus |
|---|---|---|
| AtMAN1 | *Arabidopsis thaliana* | AT1G02310 |
| AtMAN2 | *A. thaliana* | AT2G20680 |
| AtMAN3 | *A. thaliana* | AT3G10890 |
| AtMAN5 | *A. thaliana* | AT4G28320 |
| AtMAN6 | *A. thaliana* | AT5G01930 |
| AtMAN7 | *A. thaliana* | AT5G66460 |
| Amt2 | *Amborella trichopoda* | XP_020532145.1 |
| Amt5 | *A. trichopoda* | XP_006836841.1 |
| Amt6-1 | *A. trichopoda* | XP_011622682.1 |
| Amt6-2 | *A. trichopoda* | XP_020521243.1 |
| Amt6-3 | *A. trichopoda* | XP_006839096.1 |
| Amt7-1 | *A. trichopoda* | XP_011623587.1 |
| Amt7-2 | *A. trichopoda* | XP_006841527.1 |
| Amt7-3 | *A. trichopoda* | XP_006841534.1 |
| Lj2-1 | *Lotus japonicus* | XP_057455511.1 |
| Lj2-4 | *L. japonicus* | XP_057455656.1 |
| Lj4-1 | *L. japonicus* | XP_057439076.1 |
| Lj4-2 | *L. japonicus* | XP_057423754.1 |
| Lj5 | *L. japonicus* | XP_057453226.1 |
| Lj6-1 | *L. japonicus* | XP_057435418.1 |
| Lj6-2 | *L. japonicus* | XP_057435419.1 |
| Lj6-3 | *L. japonicus* | XP_057458645.1 |
| Lj7-1 | *L. japonicus* | XP_057449353.1 |
| Lj7-2 | *L. japonicus* | XP_057434964.1 |
| Os1 | *Oryza sativa* | NP_001405905.1 |
| Os2 | *O. sativa* | NP_001388559.1 |
| Os3 | *O. sativa* | XP_015629243.2 |
| OsMAN5 | *O. sativa* | XP_015638320.1 |
| Os6 | *O. sativa* | NP_001408517.1 |
| Os7 | *O. sativa* | NP_001410065.1 |
| Os8 | *O. sativa* | XP_015620164.1 |
| Os9 | *O. sativa* | XP_015624964.1 |
| PtrMAN6 | *Populus trichocarpa* | AFX59322.1 |
| Sm1 | *Selaginella moellendorffii* | XP_024527969.1 |
| Sm2 | *S. moellendorffii* | XP_002985744.2 |
| Sm3 | *S. moellendorffii* | XP_024515267.1 |
| Pp1 | *Physcomitrium patens* | XP_024377758.1 |
| Pp2-1 | *P. patens* | XP_024377738.1 |
| Pp2-2 | *P. patens* | XP_024375487.1 |
| Pp4 | *P. patens* | XP_024392427.1 |
| Pp5 | *P. patens* | XP_024373004.1 |
| Pp7-1 | *P. patens* | XP_024379491.1 |
| Pp7-2 | *P. patens* | XP_024374910.1 |
| Pp9 | *P. patens* | XP_024359301.1 |
| Mp1 | *Marchantia polymorpha* | PTQ45960.1 |
| Mp2 | *M. polymorpha* | PTQ31524.1 |
| Mp3 | *M. polymorpha* | PTQ31526.1 |
| AnMAN5A | *Aspergillus nidulans* | ABF50863.1 |
