## Supplementary material for "Atypical endo-β-1,4-mannanases are necessary for normal glucomannan synthesis in Arabidopsis": Sup. Figs.

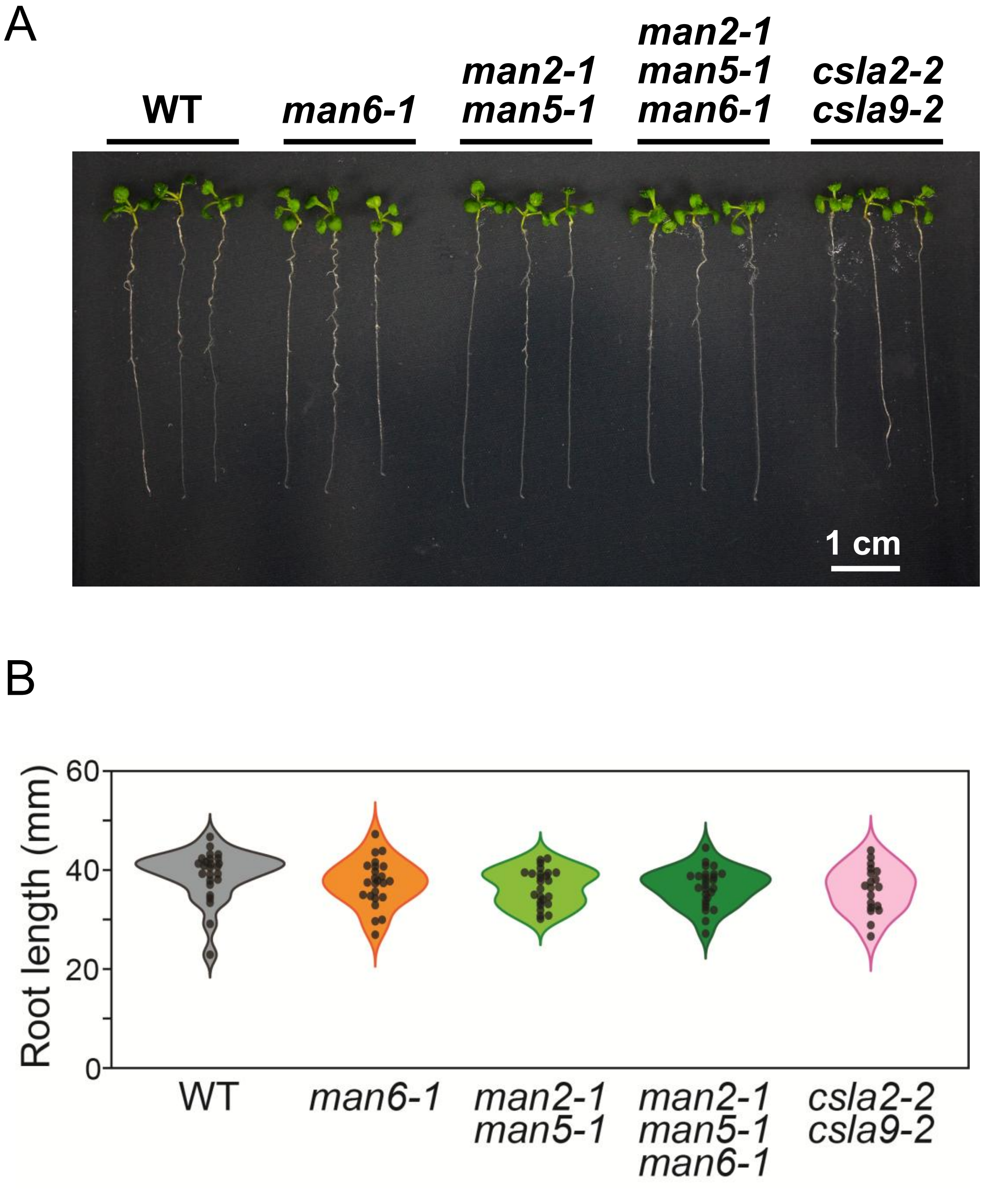


**Supplementary Fig S1. Growth phenotypes of seedlings of *man* mutants.** (A) Arabidopsis *man* mutants grown on MS medium for 9 days. (B) Root length of *man* mutants. Representative plants are shown. The seedlings of *man* mutants did not show any significant growth defects. Data are mean values with standard deviation (WT, n = 25; *man6-1*, n = 24; *man2-1 man5-1*, n = 25; *man2-1 man5-1 man6-1*, n = 27; *csla2-2 csla9-2*, n = 19). Significant differences were not found in the Tukey-Kramer test ($p < 0.05$).

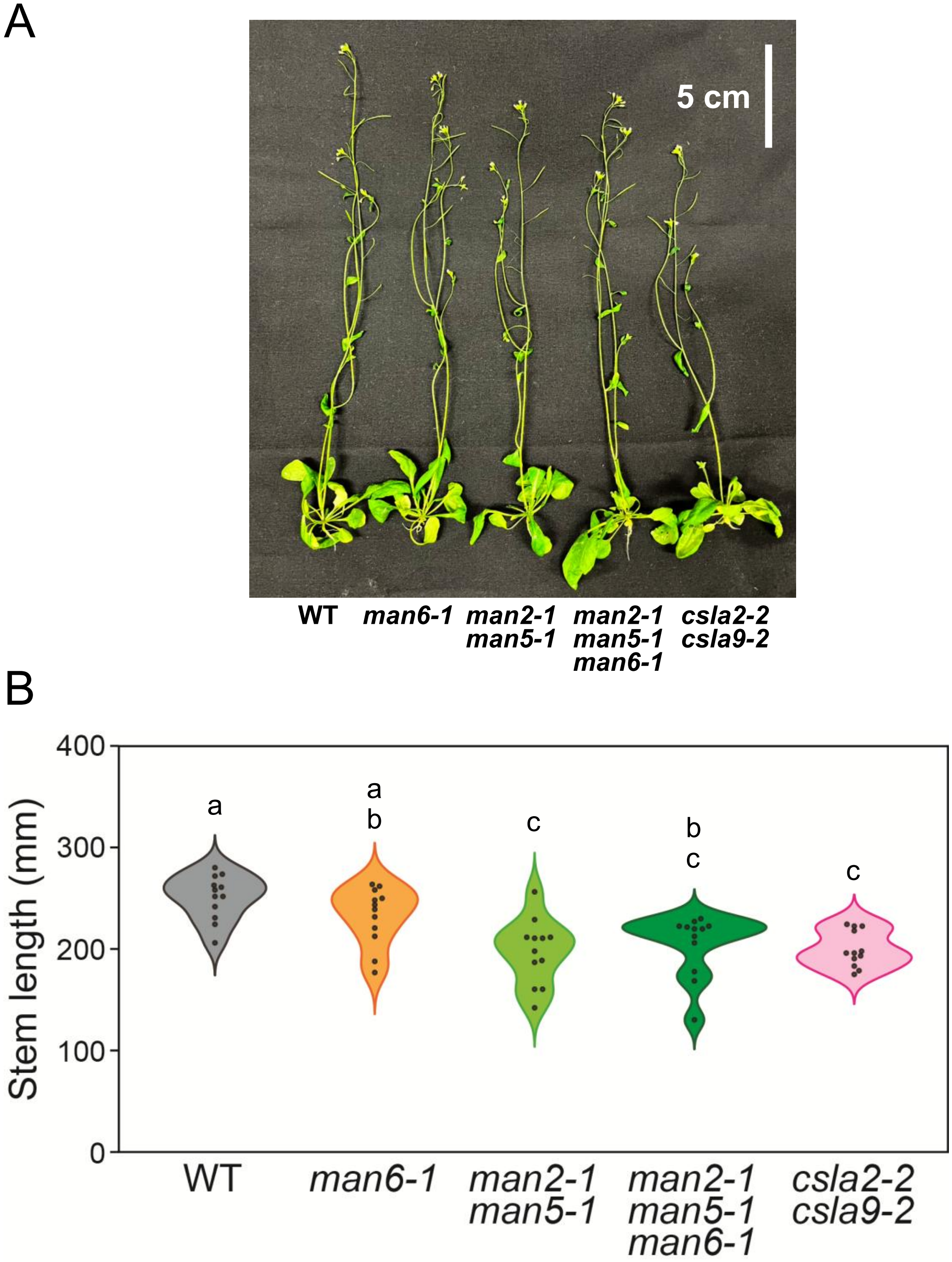


**Supplementary Fig S2. Growth phenotypes in the inflorescence stems of *man* mutants.** (A) Arabidopsis *man* mutants grown on MS medium for 12 days and then on rockwool fiber for 4 weeks. Representative plants are shown. (B) The length of inflorescence stems of *man* mutants. The *man2-1 man5-1* double *and man2-1 man5-1 man6-1* triple mutants had slightly shorter stems as observed for *csla2-2 csla9-2* double mutant. Data are mean values with standard deviation (n = 12). Different letters indicate significant differences determined with the Tukey HSD test ($p < 0.05$).

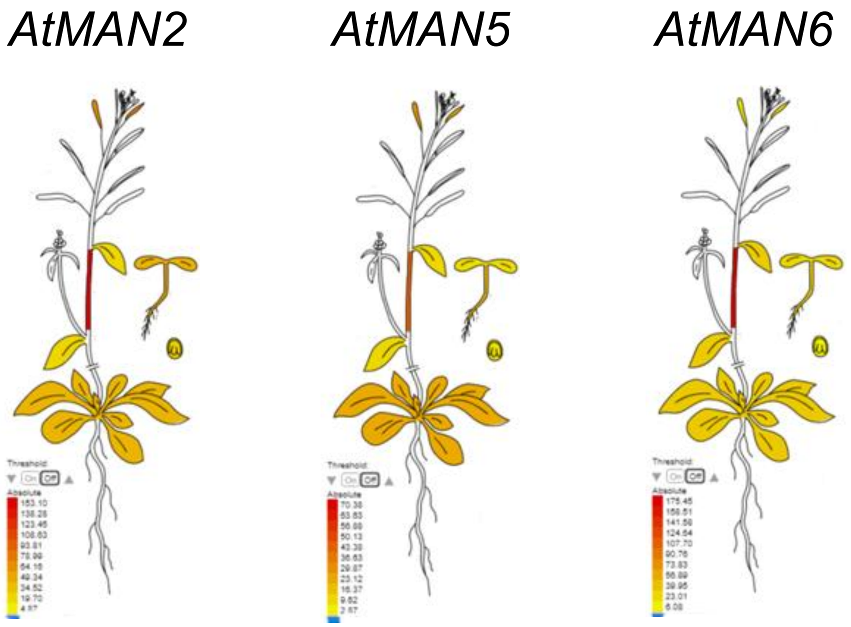


**Supplementary Fig S3. Expression patterns of *AtMAN2*, *AtMAN5*, and *AtMAN6* genes.** The expression patterns of *AtMAN2*, *AtMAN5*, and *AtMAN6*. The data were obtained from the Arabidopsis eFP Browser. The *AtMAN2*, *AtMAN5*, and *AtMAN6* genes exhibit relatively high expression levels in the inflorescence stem.

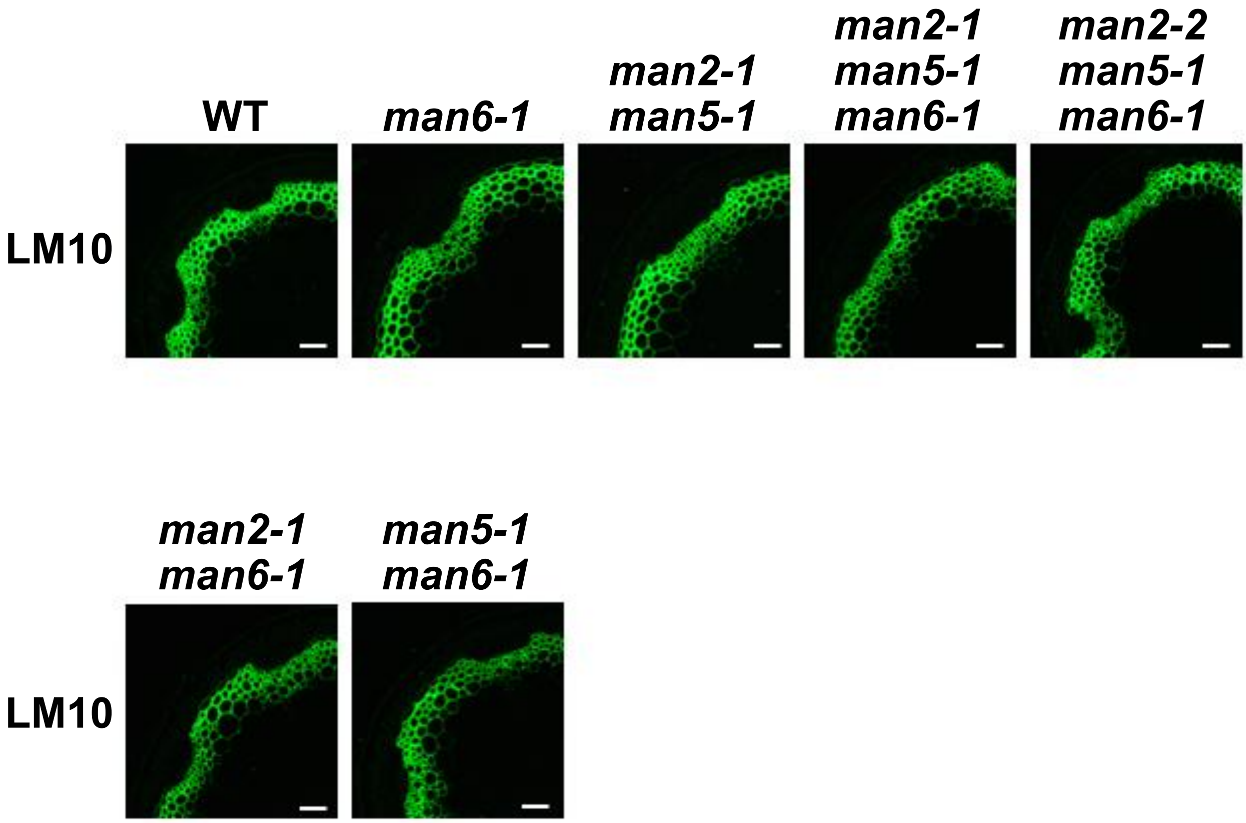


**Supplementary Fig. S4. Xylan accumulation in *man* mutants.** Cross sections from the bottom part of inflorescence stems of *man* mutants were observed. The images of xylan signals detected with LM10 antibody are shown. These *man* mutants did not show any changes in xylan accumulation. The bars indicate 50 μm.

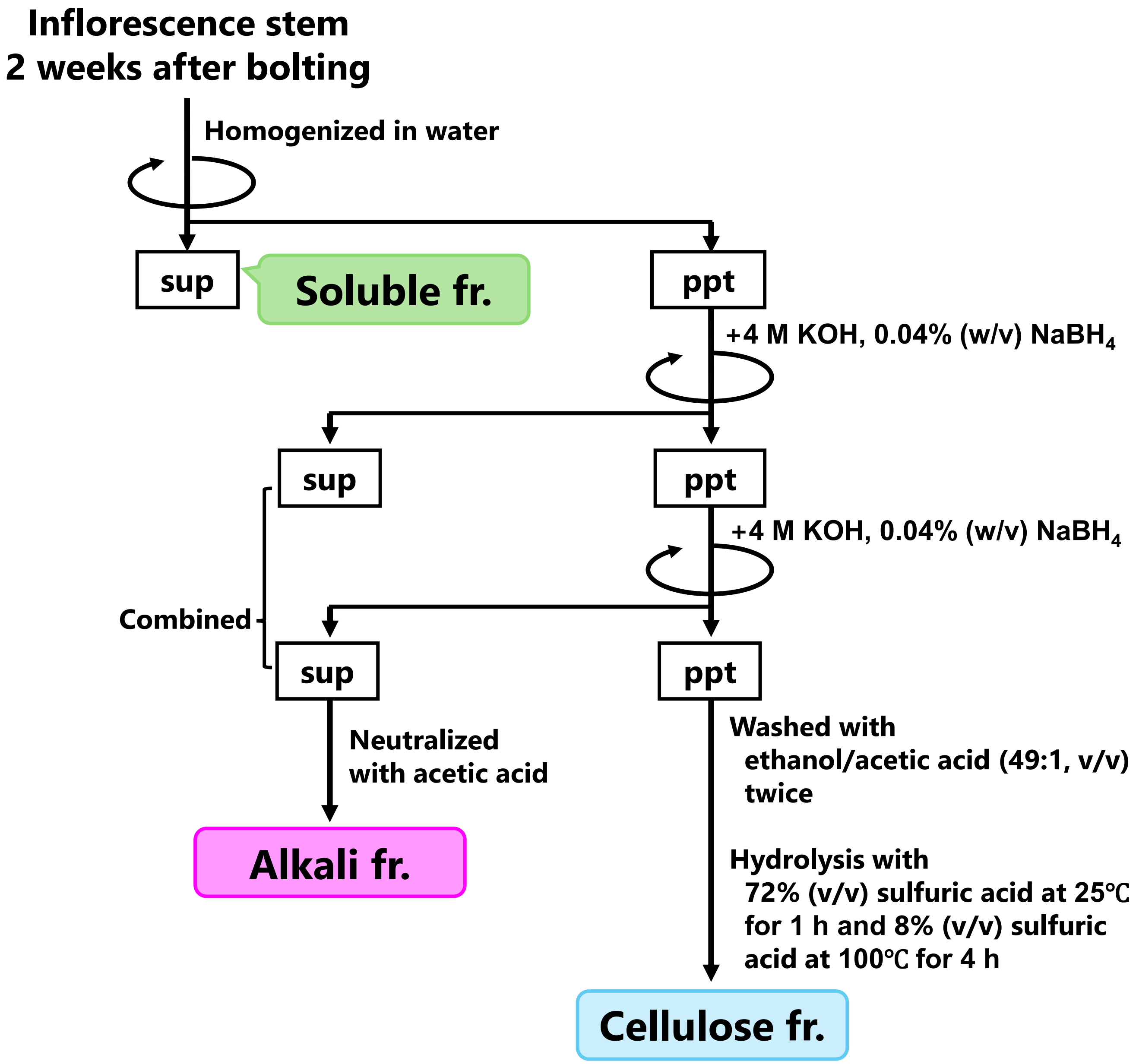


**Supplementary Fig. S5. Cell wall fractionation from inflorescence stems.** Cell wall polysaccharides were fractionated into soluble, alkali, and cellulose fractions. The sugar content and composition of each fraction were determined. fr, fraction; ppt, precipitate; sup, supernatant.

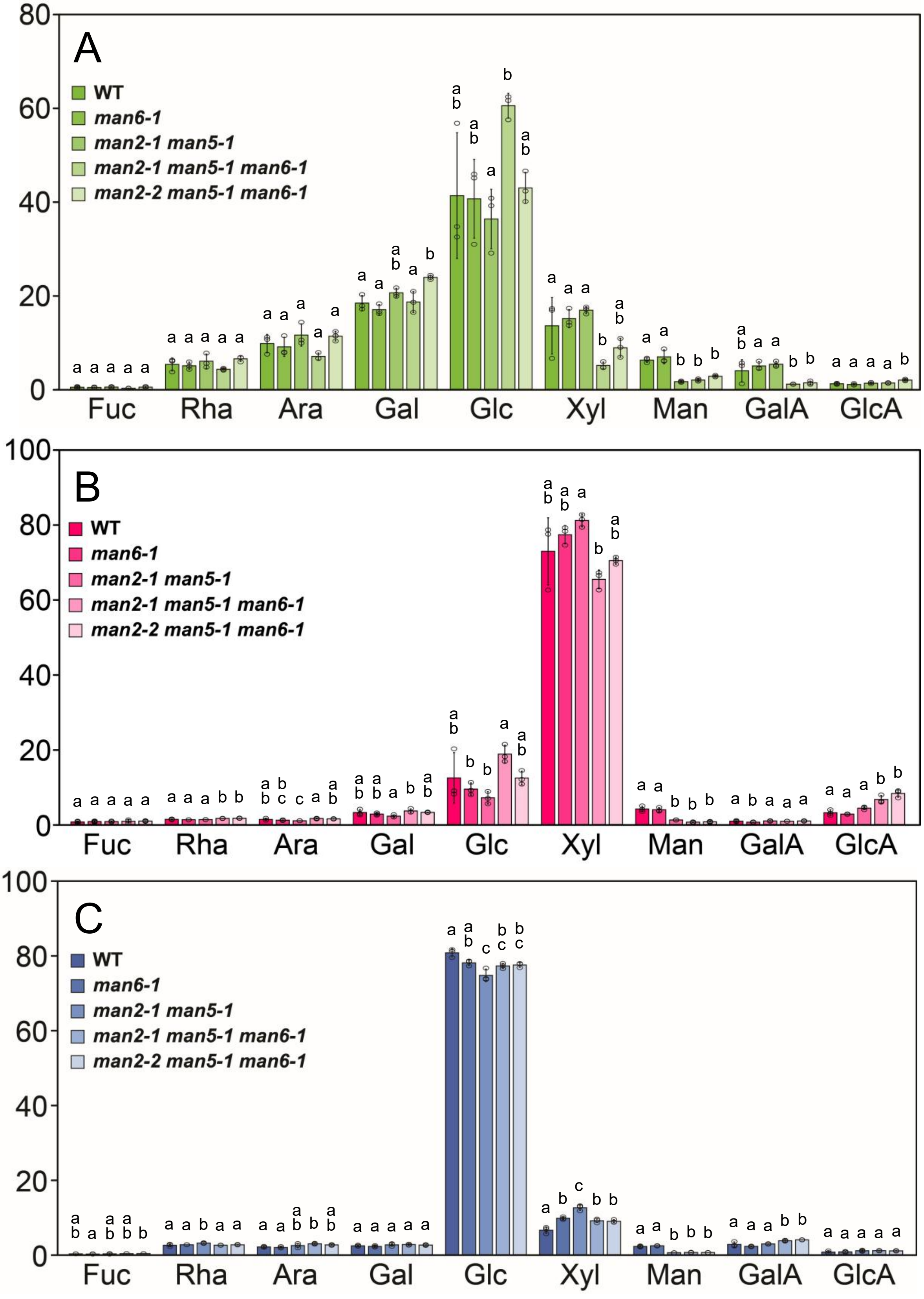


**Supplementary Fig S6. Monosaccharide composition of the soluble, alkali, and cellulose fractions.** The soluble (A) and alkali (B) fractions were hydrolyzed with 2 M TFA and the cellulose fraction (C) was hydrolyzed with 72% sulfuric acid. The monosaccharide composition was determined by HPEAC-PAD. Data are mean values with standard deviation (three biological replicates). Different letters indicate significant differences determined with the Tukey HSD test ($p < 0.05$).

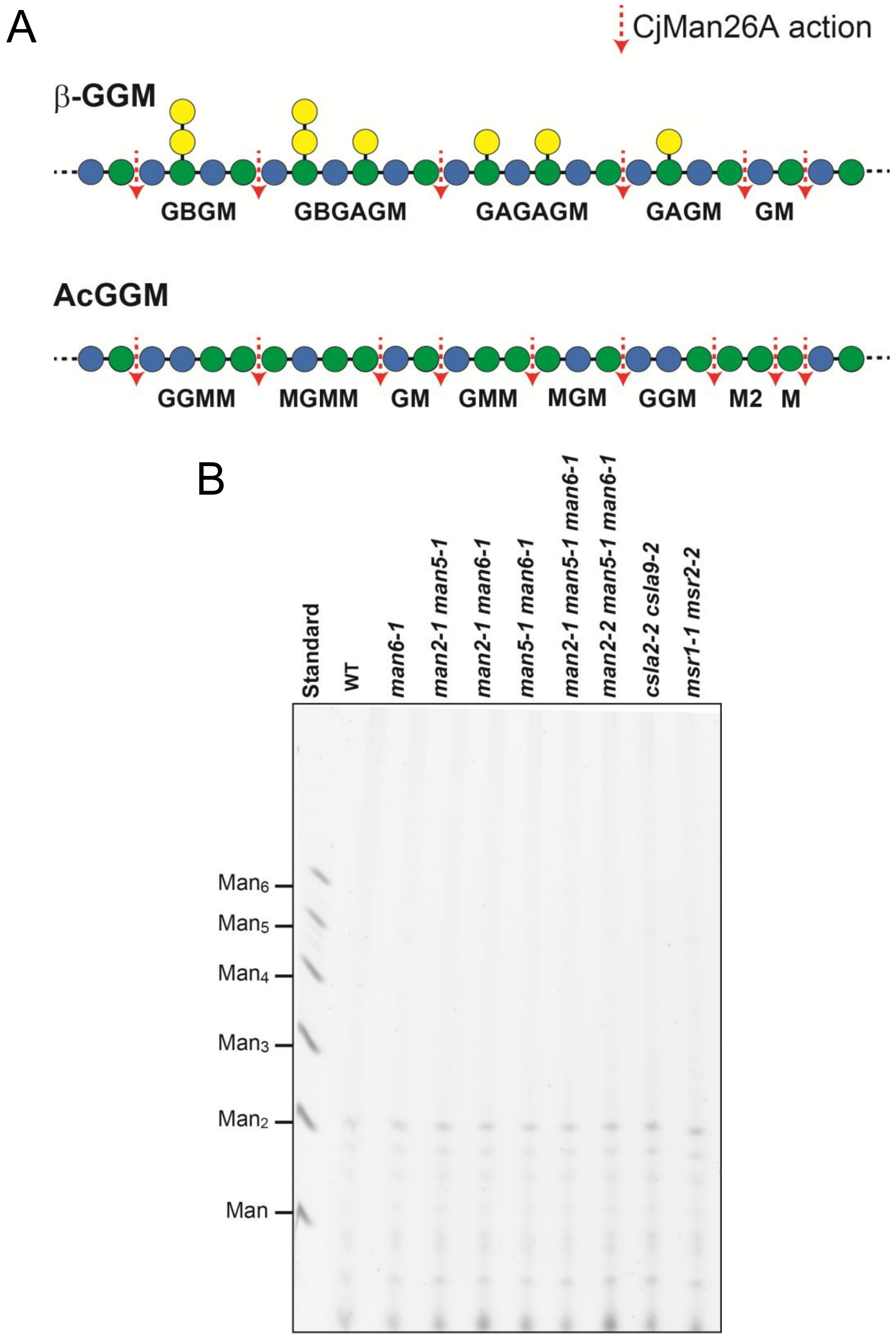


**Supplementary Fig. S7. Oligosaccharides detected by PACE.** (A) Oligosaccharides released from β-GGM and AcGGM. β-GGM highly substituted with single Gal or galactobiose gives oligosaccharides such as GAGM and GBGM. On the other hand, AcGGM mainly gives triose and biose without Gal. (B) PACE gel of control experiment. Cell wall samples without CjMan15A digestion were applied to identify non-specific bands.

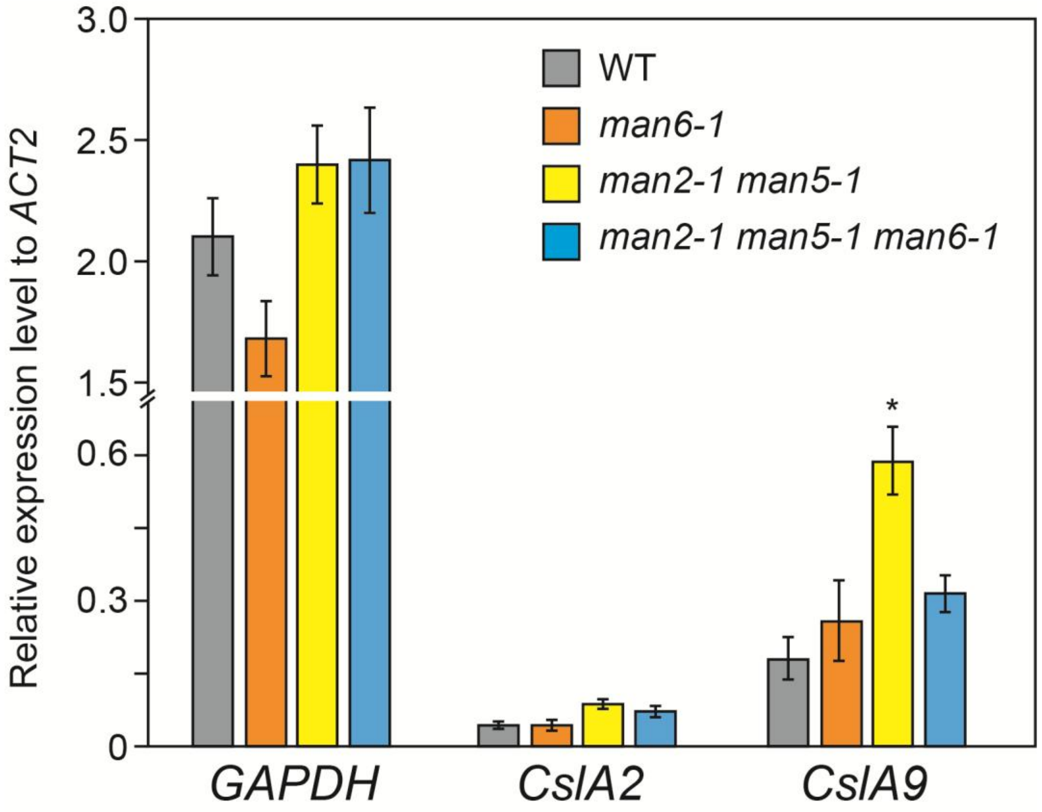


**Supplementary Fig. S8. Expression levels of *CslA2* and *CslA9* in *man* mutants.** The expression levels of *CslA2* and *CslA9* in the bottom part of inflorescence stems at 2 weeks after bolting were investigated by quantitative PCR with specific primers listed in Supplementary Table S1. Relative expression levels to *ACTIN2* (AT3G18780) are shown. Similar expression levels of a glyceraldehyde-3-phosphate dehydrogenase gene (*GAPDH*, AT3G26650) in these mutants were observed. Data are mean values with SE (5 biological replicates). The asterisk indicates a significant difference from the WT (Student's t test, *, $P < 0.05$).

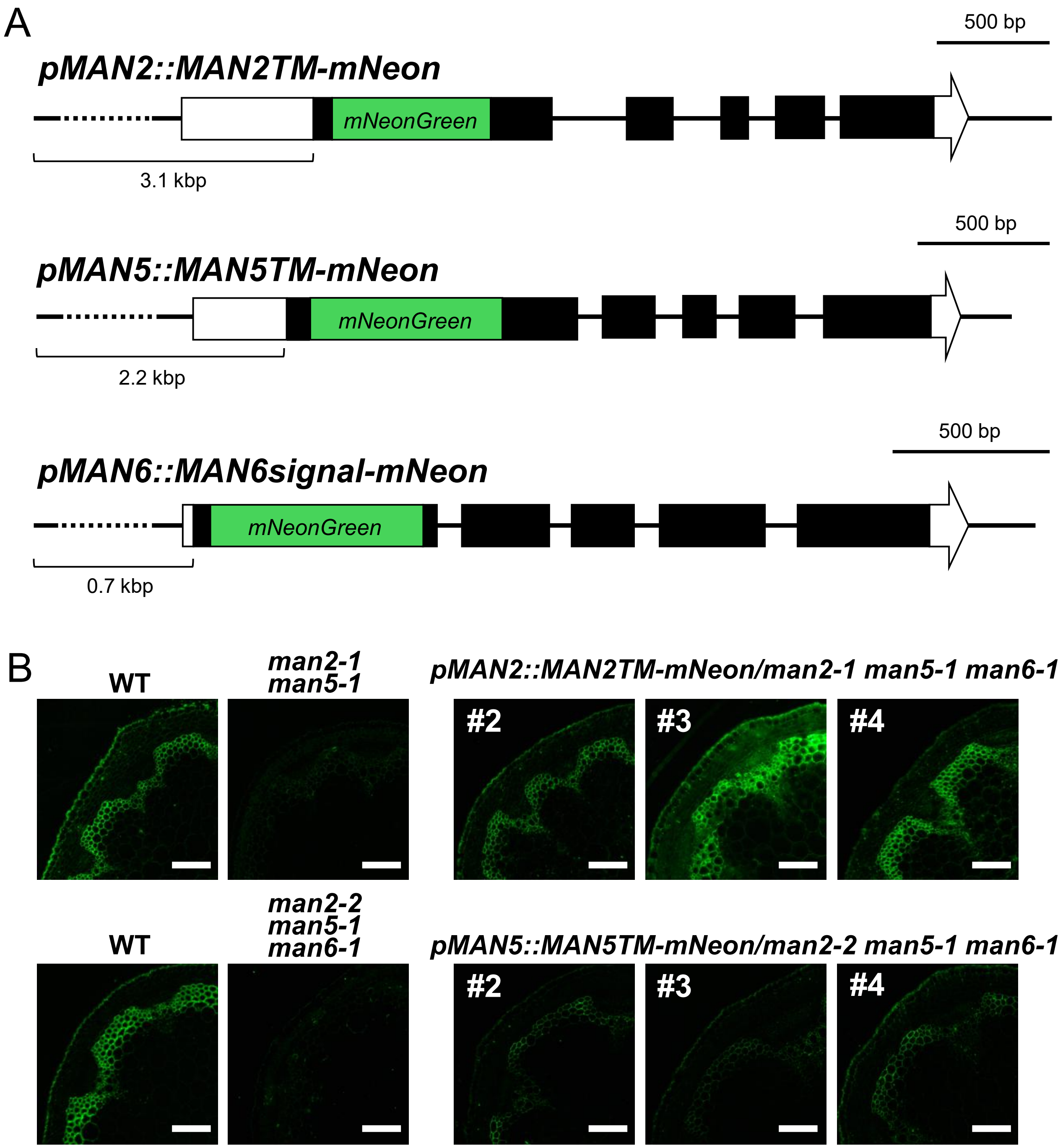


**Fig. S9. Gene constructs and complementation of *man* mutants.** (A) Schematic diagram of *pMAN2::MAN2TM-mNeon*, *pMAN5::MAN5TM-mNeon*, and *pMAN6::MAN6signal-mNeon* constructs. These constructs contained 3.1, 2.2, and 0.7 kb upstream sequences from the translation initiation codon, respectively. Black and white boxes show the coding regions and untranslated regions, respectively. (B) Gene complementation of the triple mutants with *pMAN2::MAN2TM-mNeon* (upper panels, lines #2, 3, and 4) and *pMAN5::MAN5TM-mNeon* (lower panels, lines #2, 3, and 4). The introduction of *pMAN2::MAN2TM-mNeon* and *pMAN5::MAN5TM-mNeon* at least partially restored the glucomannan accumulation in the stem of the triple mutants. For the observation of MAN-mNeon localization, *pMAN2::MAN2TM-mNeon* line #3 and *pMAN5::MAN5TM-mNeon* line #4 were used. Glucomannan was detected with LM21. The bars indicate 100 μm.

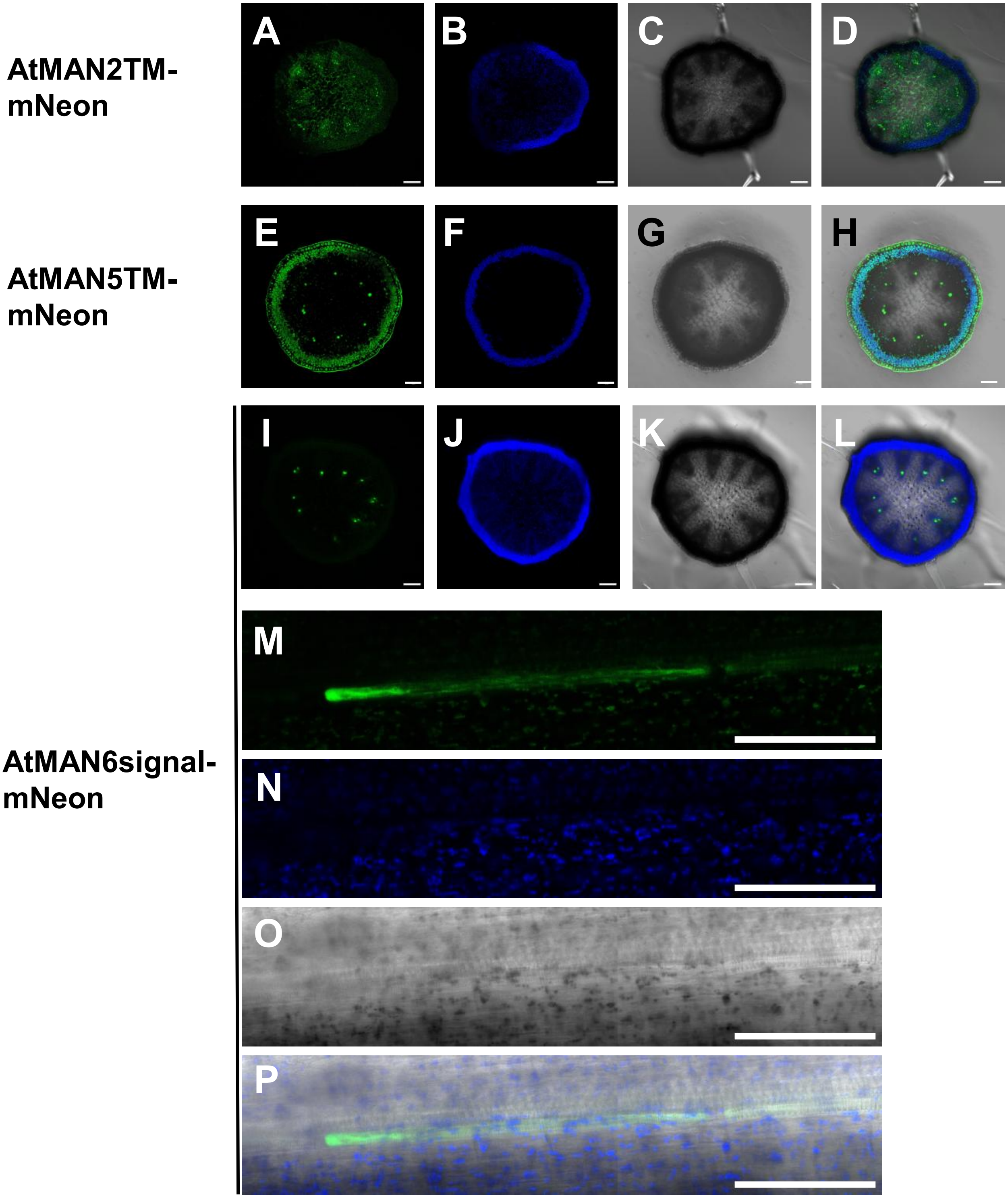


**Supplementary Fig. S10. Tissue localization of MAN-mNeon proteins.** Cross-sections of the top part of inflorescence stems of transgenic Arabidopsis harboring *pMAN2::MAN2TM-mNeon* line #3 (A-D), *pMAN5::MAN5TM-mNeon* line #4 (E-H), and *pMAN6::MAN6signal-mNeon* line #3 (I-P) were observed. Images for mNeonGreen (A, E, I, M), autofluorescence (B, F, J, N), and transmitted photomultiplier tube (C, G, K, O) were obtained and merged (D, H, L, P). Longitudinal section images of *pMAN6::MAN6signal-mNeon* line #3 (M-P) are also shown. The bars indicate 100 µm. The gene constructs are shown in Supplementary Fig. S9.

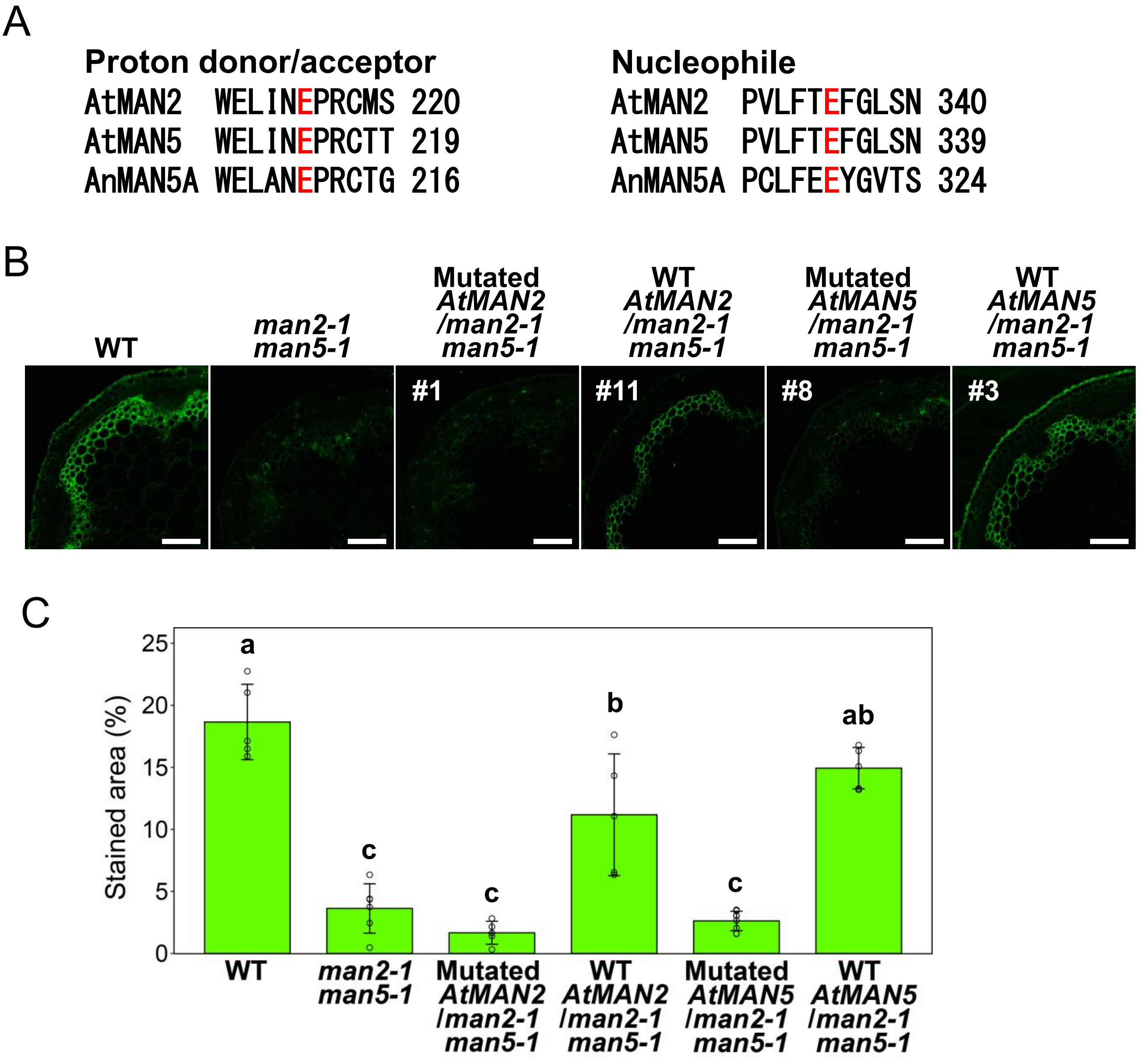


**Fig. S11. Gene complementation experiment with mutated genomic *AtMAN2* gene.** (A) Putative proton donor/acceptor and nucleophile residues in MANs. The regions including catalytic residues of AtMAN2 and AtMAN5 are aligned with AnMAN5A in *A. nidulans* (ABF50863.1). The catalytic residues are shown in red. Point mutations (E215A and E335A for AtMAN2; E214A and E334A for AtMAN5) were introduced at these catalytic residues. (B) Glucomannan accumulation in the inflorescence stem of transformants. The *man2-1 man5-1* double mutant was transformed with the mutated and WT genomic *AtMAN2* and the mutated and WT genomic *AtMAN5* genes. The representative T1 generation plants (mutated *AtMAN2/man2-1 man5-1* line #1; WT *AtMAN2/man2-1 man5-1* line #11; mutated *AtMAN5/man2-1 man5-1* line #8; WT *AtMAN5/man2-1 man5-1* line #3) are shown. Glucomannan was detected with LM21. The bars indicate 100 µm. (C) Glucomannan-detected area. Area stained with LM21 in the cross section was quantified using Fiji software. Data are mean values with standard deviation (WT, n = 5; *man2-1 man5-1*, n = 6; mutated *AtMAN2/man2-1 man5-1*, n = 5; WT *AtMAN2/man2-1 man5-1*, n = 5; mutated *AtMAN5/man2-1 man5-1*, n = 8; WT *AtMAN5/man2-1 man5-1*, n = 5). Different letters indicate significant differences determined with the Tukey-Kramer test ($p < 0.05$).

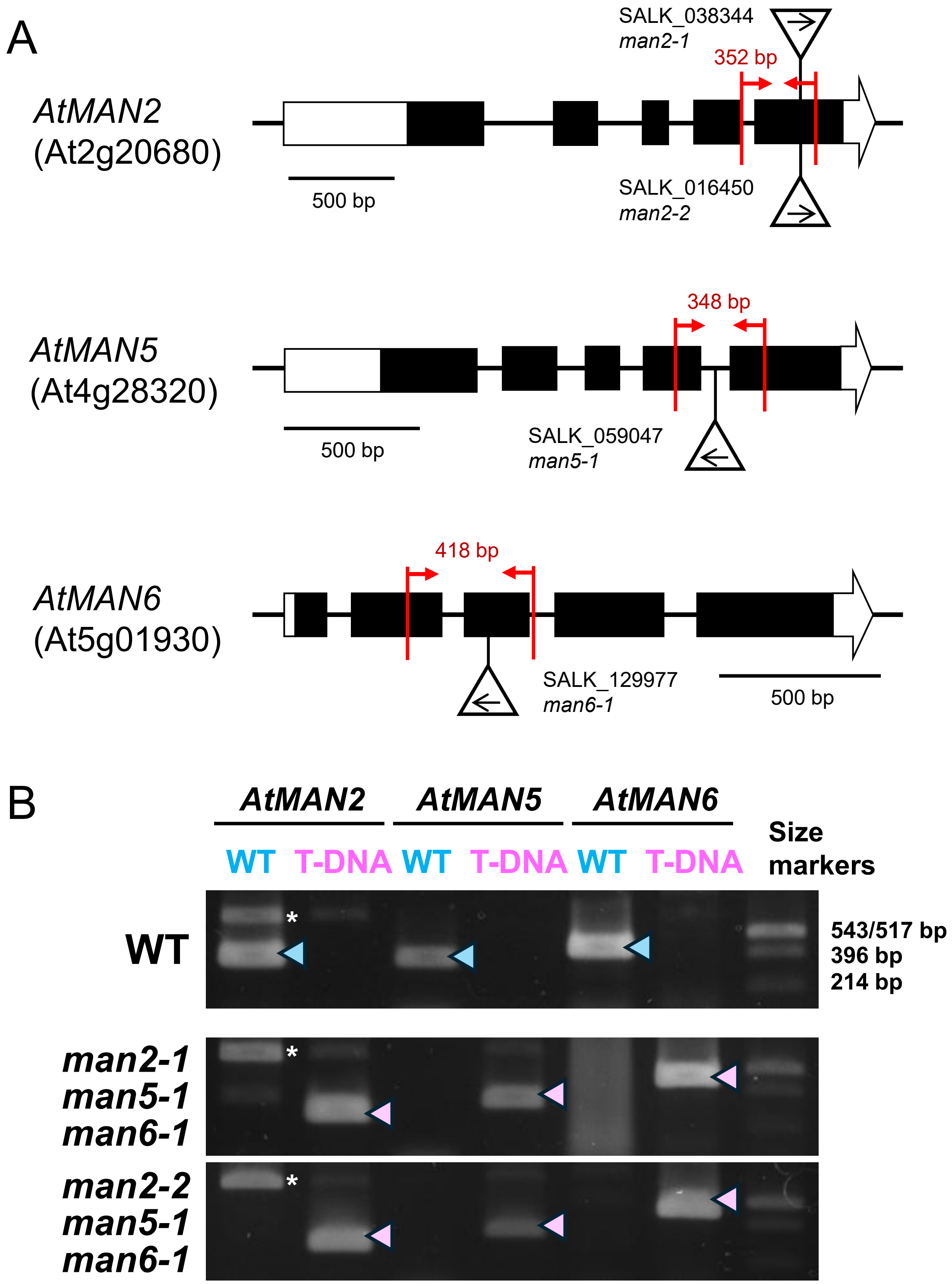


**Supplementary Fig. S12. T-DNA insertion sites and genotyping of *man* mutants.** (A) Schematic diagrams for the T-DNA insertion sites in *man* mutants. Black and white boxes show the coding and untranslated regions, respectively. Arrows represent primers used for genotype determination, which are listed in Supplementary Table S1. (B) Representative results of genotyping. The genotypes of WT and the *man2-1 man5-1 man6-1*, and *man2-2 man5-1 man6-1* triple mutants were determined by genomic PCR. The arrowheads indicate amplified DNA fragments. The asterisk indicates a non-specific PCR product.
